# AutoMod: A Software Tool for High-Accuracy Design of Curved DNA Origami Nanostructures

**DOI:** 10.64898/2026.10.20.760446

**Authors:** Elina Heikkilä, Abraham Kipnis, Taru Lautamatti, Ashwin Karthick Natarajan, Anton Kuzyk

## Abstract

In the design of DNA origami (DO) nanostructures, structural information, such as equilibrium conformation, is encoded by nucleotide sequences. The problem of finding the correct sequences for a certain target conformation is a demanding computational task, commonly addressed with the help of computer-aided design software. However, strongly curved structures where DNA helices span non-closed routes have remained extremely challenging to realize. Here, we present AutoMod, a software tool that attains significantly increased correspondence of an ideal target shape with the semi-automatically generated design. Dissimilarity of a target with a design is measured with root mean square deviation between simplified point set representations of the target shape and the computationally predicted equilibrium conformation of a design. Compared to previous methods, statistically significant decreases in mean root mean square deviation are observed with all tested target shapes. A benchmark DO structure, designed with AutoMod, is realized experimentally and its conformation is resolved with single-particle reconstruction. Subsequently, high agreement with computational results is observed. We anticipate improvements in design accuracy to facilitate the successful application of DO nanostructures in fields such as nanofabrication, biomedical engineering, and biosensing.

## Introduction

Since its inception in the 1980s [1, 2], structural DNA nanotechnology has allowed the programmable design of various nanoscale structures, including polyhedra and wireframe assemblies [3, 4, 5, 6, 7, 8, 9, 10, 11, 12], 2D [13, 14, 15, 16, 17] and 3D arrays [18, 19], concentric rings and curved shells [20, 21], bundles [22], tubes [23], boxes [24], and many other lattice-based 3D objects [25, 26, 27, 28]. These structures can self-assemble from constituent DNA strands in a highly predictable manner due to the energetic favourability of hybridization reactions between strands with complementary sequences [29]. Once formed, their shape is maintained by a certain type of immobile Holliday junctions, i.e., antiparallel crossovers [30], between DNA helices.

Currently, one of the most widely used techniques for DNA nanostructure design is the DNA origami (DO) technique [31, 32], in which a set of short “staple” strands are designed with sequences that are complementary to segments of a long circular “scaffold” strand. Commonly, the sequence of one staple contains portions complementary to at least two nonadjacent regions of the scaffold. As the staples hybridize with the scaffold, the covalent bonds in the sugar-phosphate backbones of the staples act as links that force nonadjacent scaffold regions to lie close to one another for the maximum number of Watson-Crick basepairs to occur between the staples and the scaffold. With this principle, staple sets can be designed to “fold” the scaffold into rationally designed shapes, and the task of designing a DNA nanostructure with a certain equilibrium conformation becomes the task of finding a suitable set of staples. With a successful staple design, DO structures self-assemble spontaneously in a mixture of constituent strands, and ideally, the equilibrium conformations of these structures correspond to the desired target shape.

Strategic placement of crossovers also allows the design of mechanical deformations that will take place in the equilibrium conformation. In other words, some helices will deviate from the standard B-form conformation for the structure as a whole to reach equilibrium. Conceptually, the basic principle is encapsulated by the idea that if the ends of two elastic beams, otherwise identical but with different lengths, are to be connected with short rigid links, the beams will have to undergo mechanical deformations (twist, bend, or stretch) before both connections can successfully take place. This principle can be applied to DO structures [20, 21, 28, 33] by treating DNA helices as elastic beams and crossovers as forced connections between them. The favourability of base pair formation can allow spontaneous structure formation, even when potential energy is stored in the elastic deformations of helices.

In early design paradigms, constant planar curvature is designed by uniformly distributing an overall length difference of Δ*l* = *θd* between helices [20, 33]. It is assumed that interhelical distance (*d*) is a constant, and helices, in their bent conformation, are circular arcs for a common central angle (*θ*) [20, 33]. Crossovers are either placed at only those locations where the backbones of participating strands are expected to be spatially as close to each other as possible [20], or with a more dense, regular spacing that forces helices to twist as well as bend [33]. Both crossover placement schemes have their own limitations, since positions with minimal twist can in some structures appear at large intervals [20], potentially leading to crossover networks that are too sparse to maintain well-defined structural shapes, whereas more dense crossover networks can result in unwanted twist deformations that manifest in the equilibrium conformation [34].

An alternative approach is to concentrate the length differences into small regions resembling corners. This method can then be used to approximate continuous curvature with linear segments connected by corner regions [28]. In practice, over the linear parts of the structure, parallel helices are linked to each other with a dense, regular crossover spacing, and length differences between helices are placed at the ends of these densely connected bundles. The length differences between the helices at the ends of the bundles, also known as edge gradients [28], are then used to control the angles between consecutive segments. This approach allows a straightforward generalization from constant planar curvature to varying curvature, not limited to a plane. However, the issue of unwanted twist deformations remains, since dense, regular crossover spacing forces helices to twist at bundle connection points, unless all length differences over bundle connections are multiples of the helical repeat.

Computer-aided design (CAD) software has been developed for the automated application of the constant, planar curvature without unwanted twist paradigm [21], and the edge gradient paradigm [28]. However, current CAD software only appear to achieve high design accuracies in certain cases, such as with enclosed structures or with certain cross sections. Sources of error include the simplifying assumptions that length differences cause only bend deformations and interhelical distance is a constant, as well as the omittance of the dependence of optimal length differences on crossover spacing, and in some cases the use of twist-inducing crossover positions. Since the design of large structures, containing thousands of nucleotides, is prohibitively time-consuming to do by hand, improving the accuracy of available CAD software is of high importance for the successful design of curved DNA structures. From the perspective of applied DNA nanotechnology, low design accuracy can render structures incapable of performing their desired functions. Areas that may benefit from improvements in the design accuracy of curved DNA nanostructures include nanoparticle synthesis [35], templated assembly [36, 37], the development of artificial membrane pores [38, 39], targeted molecular delivery systems [40, 41], and therapeutic protein-DNA [42] as well as mRNA-DNA [43, 44] hybrid structures.

Here, we introduce a novel software tool, called AutoMod (*Auto*mated *Mod* ification Placement for DNA Origami Design), which has been developed to address the aforementioned issue of low design accuracy with curved structures. AutoMod aims to offer improved design accuracy for continuously curved DNA nanostructures and, consequently, to facilitate the actualization of structures that have previously remained unattainable.

To demonstrate the differences between alternative design approaches, we designed a benchmark structure with constant, planar curvature with all currently known methods and software for the design of curved multilayer lattice-based DO structures, including AutoMod, and estimated the equilibrium conformation for each one of these alternative designs with molecular dynamics simulations. We then compared the results with the target shape to quantify the accuracy of each design method and software. For a more comprehensive comparison between AutoMod and previous CAD software capable of automated design generation, including automated crossover placement and not limited to constant planar curvature, we present a gallery of designs aiming to realize target shapes with various cross-sectional profiles and containing curvatures of different magnitudes, alongside their accuracy scores. To experimentally confirm the simulated findings, we self-assembled the benchmark structure with the DO technique and analysed its equilibrium conformation with cryogenic Transmission Electron Microscopy (cryoTEM) and single-particle reconstruction.

As a result, we observe that AutoMod achieves statistically significant improvements in design accuracy for all test designs. Compared to previous methods, we measure decreases in the discrepancy between the target and the computationally predicted equilibrium conformation for the initial design, quantified with the Root Mean Square Deviation (RMSD) between simplified point set representations. We observe percentage changes between -13% and -80% for a total of 9 example designs, with an average of -56%. We find that the single-particle reconstruction of the benchmark structure displays a conformation similar to the computational prediction. In summary, this work expands the existing DO toolbox by introducing a software tool for the design of curved DO nanostructures.

## Materials and Methods

### AutoMod

AutoMod is a Python package that allows users to design continuously curved structures via a Graphical User Interface (GUI). In practice, a typical design process, as illustrated in Figure 1, would proceed as follows. First, the user decides on a target shape and a crosssectional profile. The target shape is then inputted as a sequence of points the structure should follow as closely as possible. Points can be added or removed, their coordinates can be changed, and their sequential ordering can be set via the main 3D view of the GUI. The cross-sectional profile is inputted by specifying the number and arrangement of helices in the cross-sectional plane via the cross section tool.

**Figure 1:**
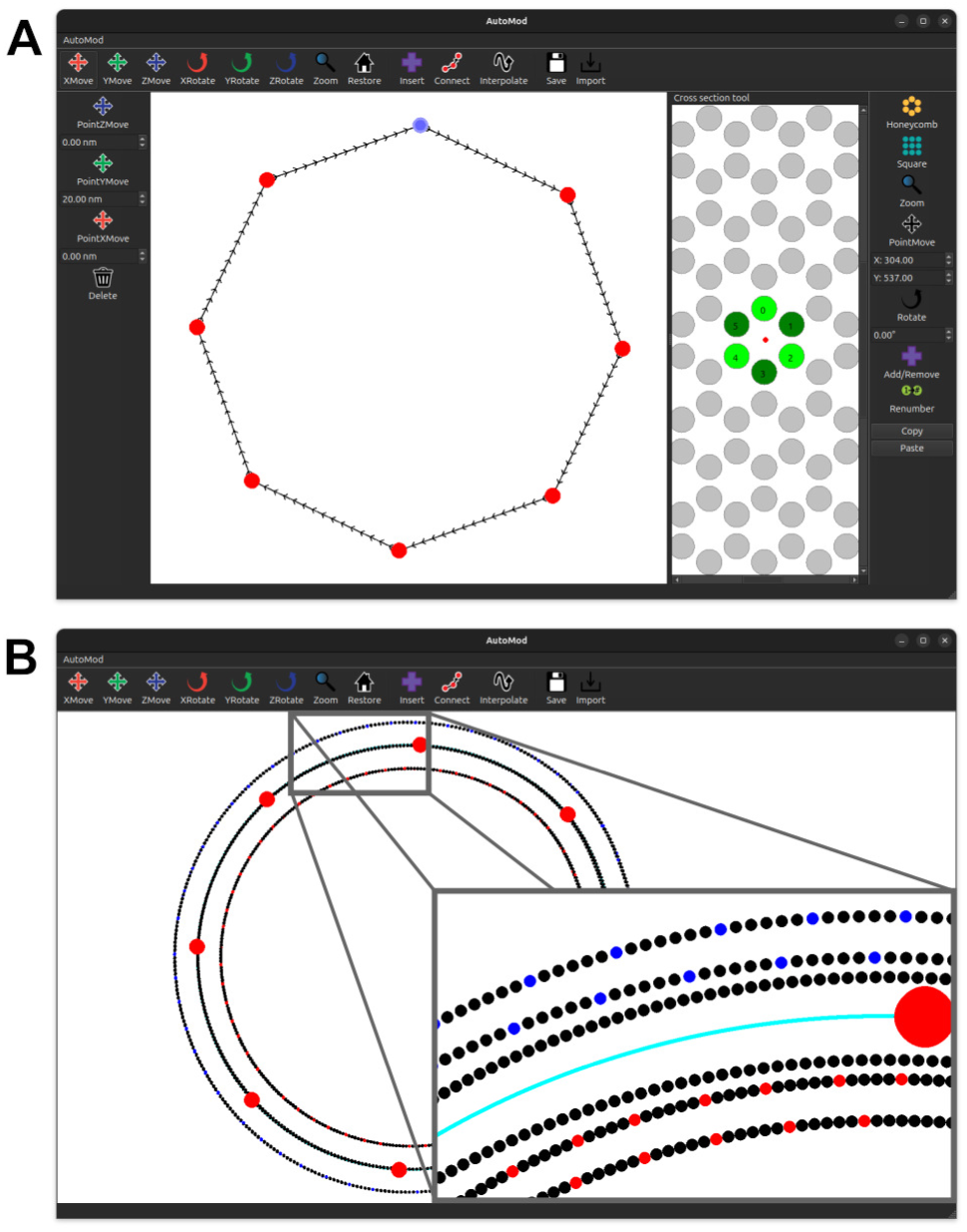
Schematic workflow of DNA origami design with AutoMod. (A) A set of target points, shown as red discs, is used to draft the target shape. A sequence of connected target points with cross-sectional profiles specifies the complete target shape. On the left, the connections are shown as black lines with arrowheads that display the direction of the connection. On the right, an exemplary cross-sectional profile containing six helices in the honeycomb lattice arrangement is shown. Helices with odd numbers are shown as dark green discs and helices with even numbers as light green discs. The parity of helix numbers is used to determine the polarities of the scaffold and staple strands. That is, in the caDNAno file format, odd helices have their scaffold strands running in the 3’-5’ direction, and even he-lices have their scaffold strands running in the 5’-3’ direction. The small red dot displays the location of the reference curve in relation to the cross-sectional profile. Here, the reference curve passes through the centre of mass. (B) A solved modification pattern after interpolation. Small black discs show the target locations for basepair centroids. Blue discs indicate locations for insertions, and red discs for deletions. The reference curve whose length is used to solve the modification pattern is shown in cyan. The part of the solution that lies within the gray rectangle is shown as enlarged and rotated in the bottom right corner.

Once the target is fully specified, a solution to the corresponding design problem can be generated with the interpolate and save functions. The interpolate function constructs target curves for helical axes as natural cubic splines. The length differences between helices in the target conformation are solved by comparing the arc lengths of the representative splines. These arc length differences are then discretized (Supplementary Equations 1) and stored in caDNAno format as modifications, i.e., insertions (addition of a base pair compared to a reference number of base pairs) and deletions (removal of a base pair compared to the reference).

The user can visually inspect the solution in the main 3D view, and alter the user settings or tune the target shape if necessary. The cross-sectional profile can be freely rotated at each point in the sequence of target points to impose minimal restrictions on the set of possible targets. Once a satisfactory solution is found, it can be written to a JavaScript Object Notation (JSON) file by saving. The JSON file format is used to ensure maximal compatibility with other software such as caDNAno [45]. At the time of saving, a crossover pattern accompanying the modification pattern is solved (Supplementary Note 1) by comparing the rotational phases of neighboring helices at potential crossover locations.

Compared to previous design software, novel features of AutoMod include the first automated implementation of the targeted modifications design method [33] for any continuously curved target shape, and the first automated solution method that recognizes the interdependence of the two parts of the full solution, i.e., the optimal modification pattern and the optimal crossover pattern, on each other.

Modifications and crossovers cannot be solved independently of each other because the lengths of the helical segments into which modifications are placed modulate the effectiveness of these modifications in inducing the desired global deformations, such as bending or twisting of the entire structure [46]. This is due to the strain energy of a deformed double-helical segment depending on its stiffness, which is, in turn, inversely proportional to its total length [47]. In other words, modifications placed into short helical segments are more effective in inducing the desired global deformations than modifications placed into longer segments [46]. In addition, the amount of error caused by the simplifying approximation that the interhelical distance is a constant throughout the structure depends on the density of the crossover pattern. This is because the distance between crossovers affects how far apart the helices will splay due to the electrostatic repulsion between their negatively charged backbones [48, 49].

In practice, if the modification pattern is kept constant, but the total number of crossovers in the modified region of the structure is increased, the magnitude of observed global curvature increases (Supplementary Figure S1). For simplicity, this relation between modifications and crossovers is often omitted in the design phase. This allows straightforward solutions at the cost of design accuracy.

AutoMod offers an alternative approach where the aforementioned dependence is acknowledged while still keeping the solution method as simple as possible. Briefly, this is done by introducing two new parameters; the effective change in the length of a helix induced by the addition or removal of a base pair (*δ*), and the target value for the average crossover density (*ρ*) of each helix. Here, average crossover density refers to the total number of crossovers relative to the total number of base pairs in a helix. Both parameters are assigned a constant value prior to solving the modification and crossover patterns. Contrary to previous design paradigms, the value of *δ* can differ from the assumed rise along the helical axis per base pair, which for B-form DNA helices is typically assumed to be approximately 0.34 nm [50]. This allows pre-emptive cancellation of errors by choosing such a pair of values that the additional length differences resulting from modification placement with *δ <* 0.34 nm approximately match the loss of global curvature due to limited crossover density.

The task of finding a suitable pair of parameter values can then be solved as a two-dimensional optimization problem. Here, a rudimentary grid search is performed to calibrate the default setting (Supplementary Figures S2-3), but both parameters can be freely adjusted via user settings, allowing structure-specific optimization and the application of more sophisticated optimization methods.

Once the parameter values are known, they can be applied to solve modifications and crossovers in a sequential manner. First, modifications are solved by assuming that their effectiveness is given by *δ*. Then, the average crossover density of each helix is followed during crossover placement and matched to the predetermined target value given by *ρ* (Supplementary Note 1). In this way, the choice of treating *δ* and *ρ* as predetermined constants allows for a simple solution, where modifications can be solved with just one pass through the structure. If the target value for the average crossover density cannot be reached for all helices, the user is notified. In this case, the target value can be lowered from the default in the user settings.

AutoMod aims to be a specialized design tool focused on solving the modification pattern needed to induce global curvature as specified by the target shape. Hence, some aspects of the design of a complete DO structure, including scaffold routing, folding kinetics, and the placement of staple nicks, are not addressed. Other software tools, such as Autobreak [51] and Scaffoldselector [52], can be utilized in conjunction with AutoMod to optimize some of the here unaddressed design choices.

### Design of DNA Origami Structures

The design process of all structures presented here follows the same series of steps, where the desired target is first specified as a combination of a parametric curve and a cross-sectional profile. The curve gives the route that the cross-sectional profile follows throughout the length of the structure, meaning that at any point on the curve, the unit tangent of the curve is the unit normal vector for the respective cross-sectional plane. The arrangement of helices in the cross-sectional plane is the same for all points on the curve and is given by the cross-sectional profile. Conventional honeycomb [25] and square lattice [26] arrangements were used to design all cross-sectional profiles.

The split ring benchmark structure, used to conduct the comparison of design accuracy across methods and software (Figure 2), was designed with the manual targeted modifications method [33] and four CAD software; ENSnano [53], DNAxiS [21], MagicDNA [28], and AutoMod. The target shape for the benchmark structure is a split ring (Figure 2A), i.e., a toroidal shape with constant curvature but disconnected ends. The radius of the target shape is *R* = 20 nm, from the midpoint of the cross section to the centre of symmetry. The cross-sectional profile contains 3 × 6 helices in a regular square lattice arrangement. With a constant interhelical distance of 0.5 nm, the cross sectional profile is 7 nm wide and 14.5 nm high.

**Figure 2:**
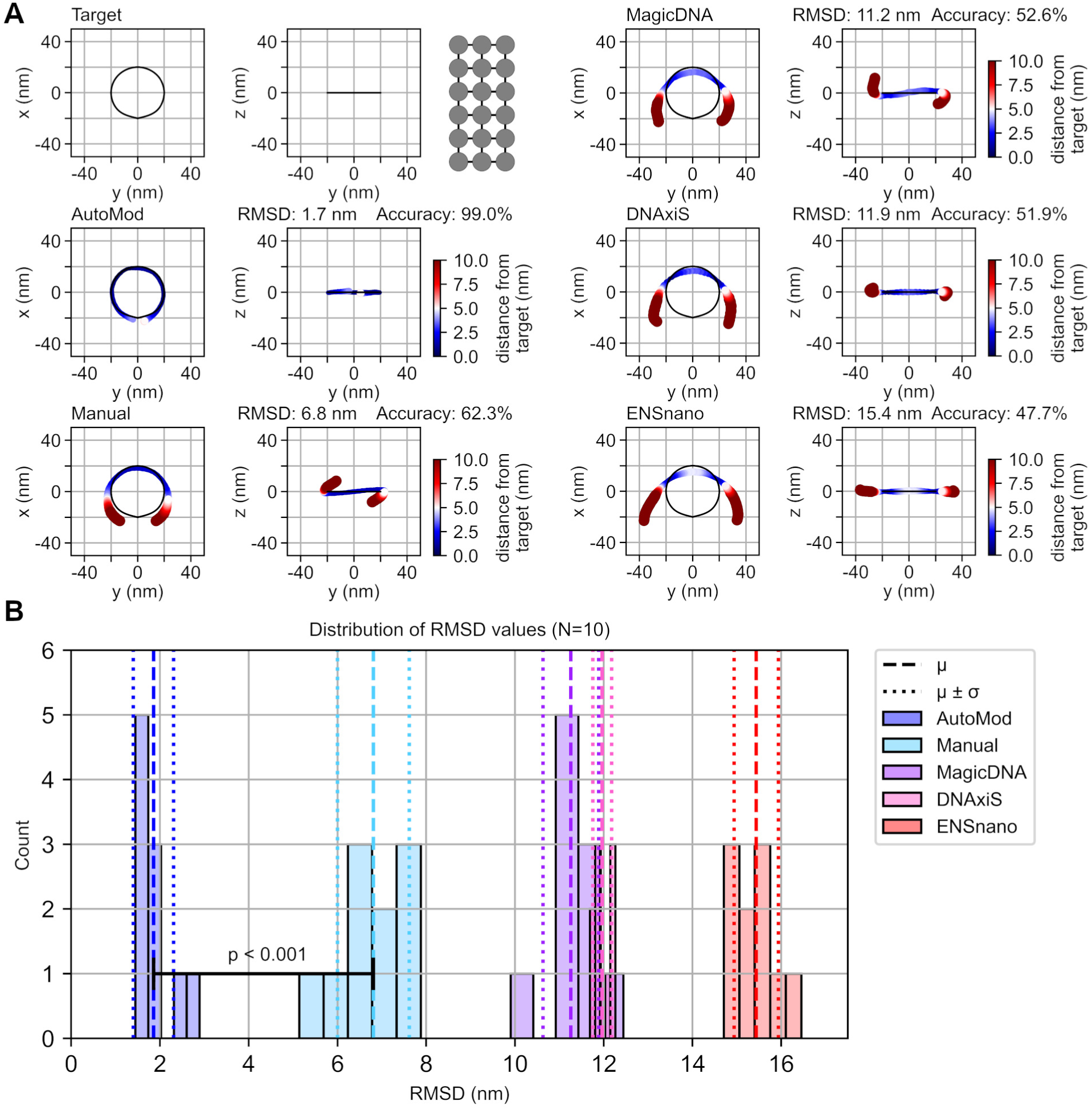
Comparison of design accuracy across methods and software with a curved bench-mark structure. (A) The point set representation of the target shape is shown along with point set representations of means from MD simulations. A distance threshold value of; *d*_lim_ = 5.0 nm, was used to compute all accuracy percentages. (B) Distributions of RMSD values for an ensemble of *N* = 10 different simulation runs for each structure. Five bins of equal width are distributed uniformly between the minimum and maximum value of each distribution. Welch’s t-test confirmed the alternative hypothesis that the mean of the AutoMod distribution is less than the second smallest mean, with a p-value of approximately 9.2 *·* 10*^−^*^11^.

This target shape was selected for benchmarking design methods and software because it is a simple shape that is, nevertheless, challenging to realize with high accuracy. Because the ends are disconnected, any error affecting the magnitude of curvature in the osculating plane, i.e., the plane in which the tangent and normal vectors of the target curve lie, or creating unwanted torsion out of the osculating plane can easily manifest itself in the equilibrium conformation. In contrast, in a complete ring with connected ends, cancelation of unwanted torsion could take place within the structure without causing clearly visible deformations in the equilibrium conformation.

The manual targeted modifications method was applied by calculating the number of modifications in each helix by hand (Supplementary Equations 2). After this, the modifications were manually added to a JSON file via the caDNAno GUI. With DNAxiS, the target shape for the benchmark structure was inputted by specifying the cross-sectional profile along its distance from the axis of revolution (Supplementary Note 2). The original output was a toroid that was turned into a split ring by nicking all strands along the same cutting plane.

For inputting the target shape into ENSnano, MagicDNA, and AutoMod, eight uniformly spaced points were sampled from the target curve (Supplementary Equations 3), and cross-sectional profiles were manually inputted via the respective GUIs. Automated crossover placement was used when available. Otherwise, manual crossover placement was used. The AutoStaple function in caDNAno was used to determine the crossover pattern for the structure with manually calculated modifications. All target shapes for the structures displayed in Figures 3 and 4 were inputted to MagicDNA and AutoMod in a similar manner by sampling a set of points from the target curve (Supplementary Equations 3) and manually inputting the cross-sectional profiles.

**Figure 3:**
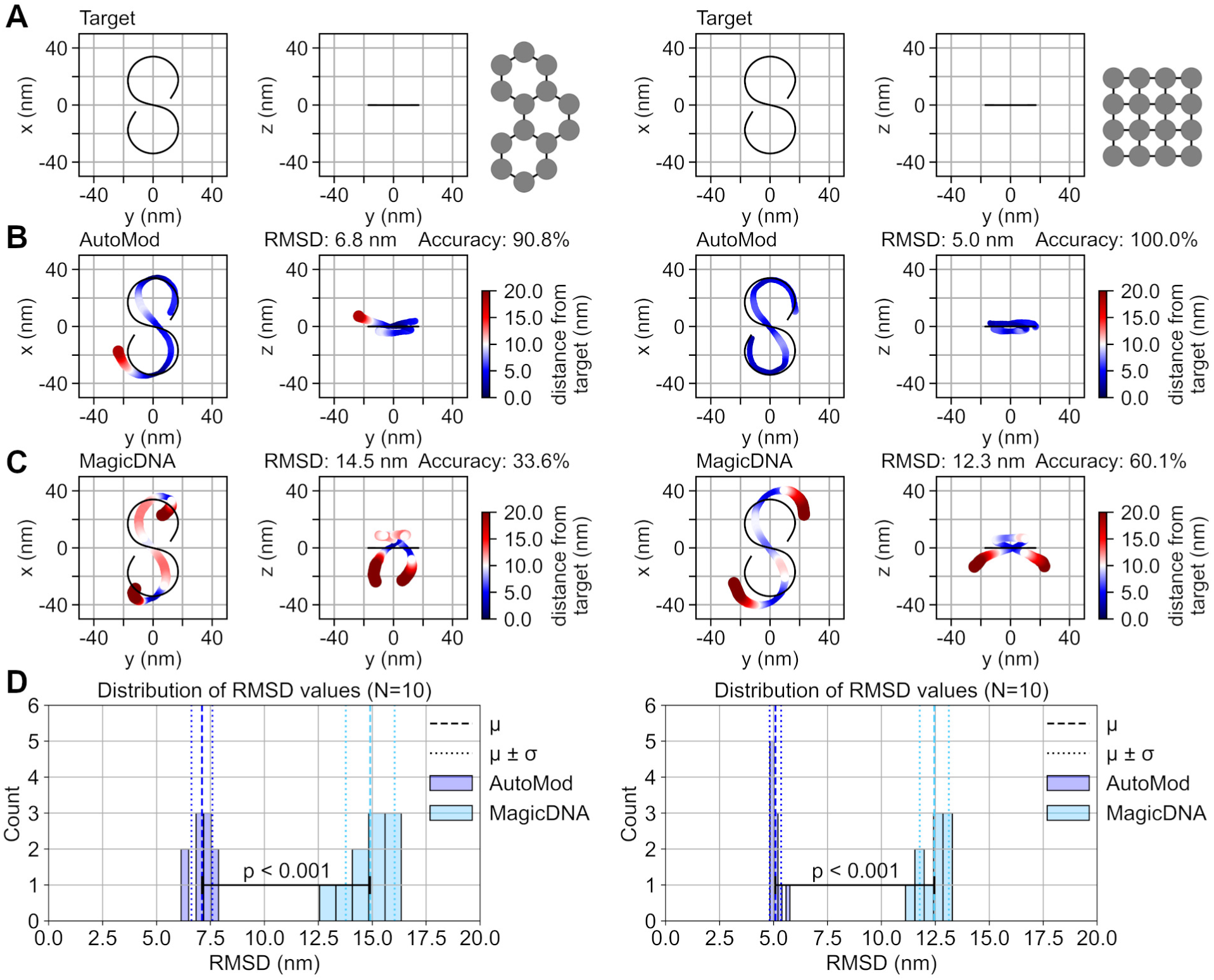
Evaluation of design accuracy with a curved target shape and two different cross-sectional profiles adhering to the honeycomb and square lattices. A distance threshold value of; *d*_lim_ = 10.0 nm, was used to compute all accuracies. (A) Point set representations of target shapes along with the cross-sectional profiles on the right. (B) Accuracy of AutoMod designs as given by the total mean over all simulation runs. (C) Accuracy of MagicDNA designs as given by the total mean over all simulation runs. (D) Distributions of RMSD values for an ensemble of *N* = 10 different simulation runs for each structure. Five bins of equal width are distributed uniformly between the minimum and maximum value of each distribution. Welch’s t-test confirmed the alternative hypothesis that the mean of the AutoMod distribution is less than the mean of the MagicDNA distribution, with p-values of approximately 9.6 *·* 10*^−^*^11^ (honeycomb lattice) and 7.8 *·* 10*^−^*^13^ (square lattice).

**Figure 4:**
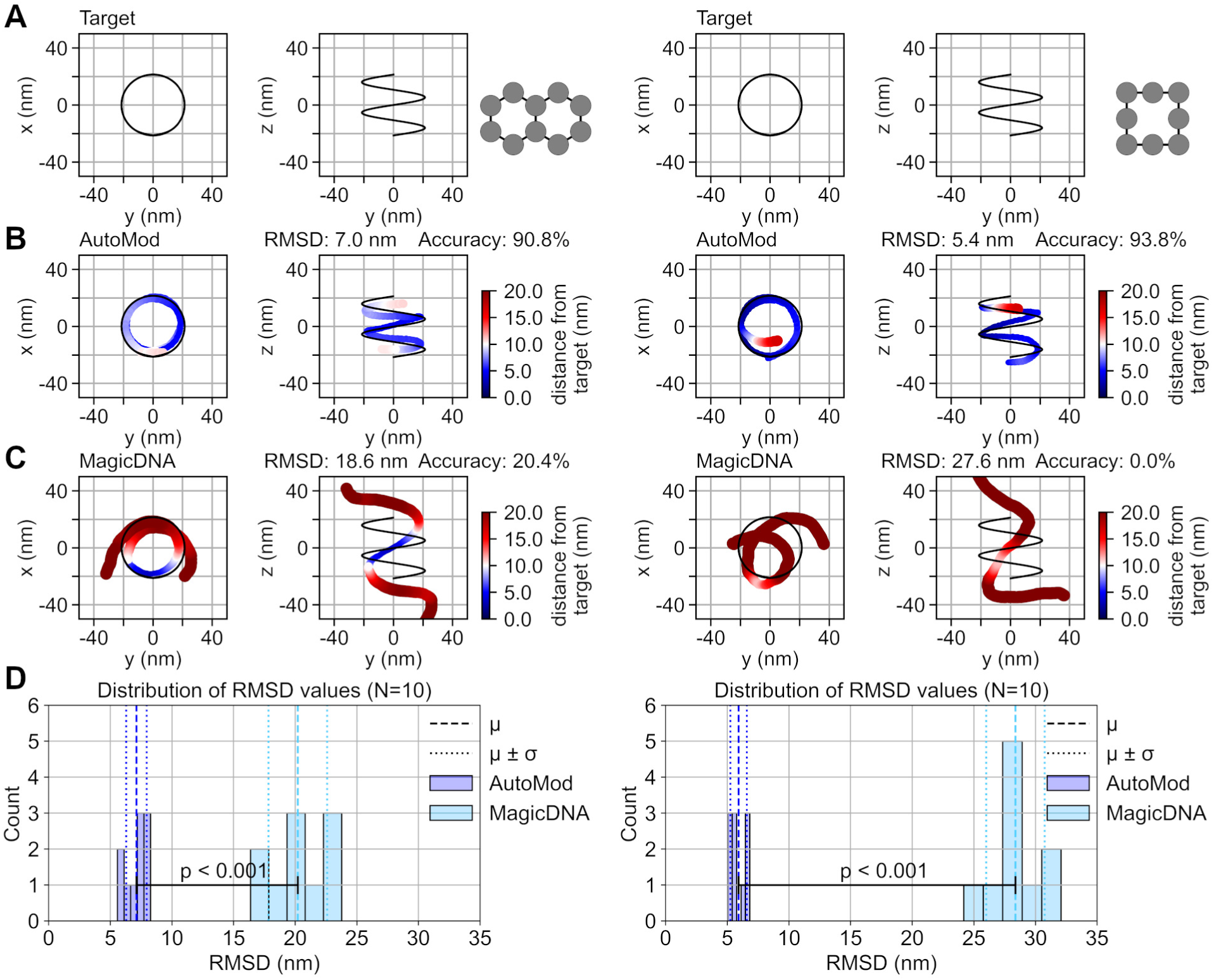
Evaluation of design accuracy with a curved target shape and two different cross-sectional profiles adhering to the honeycomb and square lattices. A distance threshold value of; *d*_lim_ = 10.0 nm, was used to compute all accuracies. (A) Point set representations of target shapes along with the cross-sectional profiles on the right. (B) Accuracy of AutoMod designs as given by the total mean over all simulation runs. (C) Accuracy of MagicDNA designs as given by the total mean over all simulation runs. (D) Distributions of RMSD values for an ensemble of *N* = 10 different simulation runs for each structure. Five bins of equal width are distributed uniformly between the minimum and maximum value of each distribution. Welch’s t-test confirmed the alternative hypothesis that the mean of the AutoMod distribution is less than the mean of the MagicDNA distribution, with p-values of approximately 3.1 *·* 10*^−^*^9^ (honeycomb lattice) and 2.4 *·* 10*^−^*^11^ (square lattice).

In MagicDNA, the extrude method was used to design the curvature inducing edge gradients for all structures. (See Supplementary Figures S4-5 for accuracy comparisons between the default extrude method, the extrude method with manually adjusted edge gradients, and the sweep method.) The extrude method was preferred over the sweep method as it resulted in a higher design accuracy in the preliminary comparison (Supplementary Figure S4). In AutoMod, the default settings (Supplementary Note 3) were used for all designs.

To minimize the effect of design choices not directly subject to the current comparison, which focuses solely on the accuracy of the induced curvature, an identical scaffold routing was used in all designs aiming to realize the same target shape, and staple breakpoints were not added to any of the designs. (See Supplementary Figure S6 for an estimate of the amount of conformational variation that can be expected due to the placement of staple breakpoints.) For example, the Autobreak tool [51] can be used in conjunction with the software compared here to find thermodynamically optimal staple breakpoints.

The experimentally realized structure (Figure 5) was designed in AutoMod with the default settings and random seed 0. (See Supplementary Figure S7 for an estimate of the amount of conformational variation that can be expected between different random seeds.) Scaffold routing was done manually in caDNAno, and staple breakpoints were determined with the Autobreak tool [51] (Supplementary Note 4).

**Figure 5:**
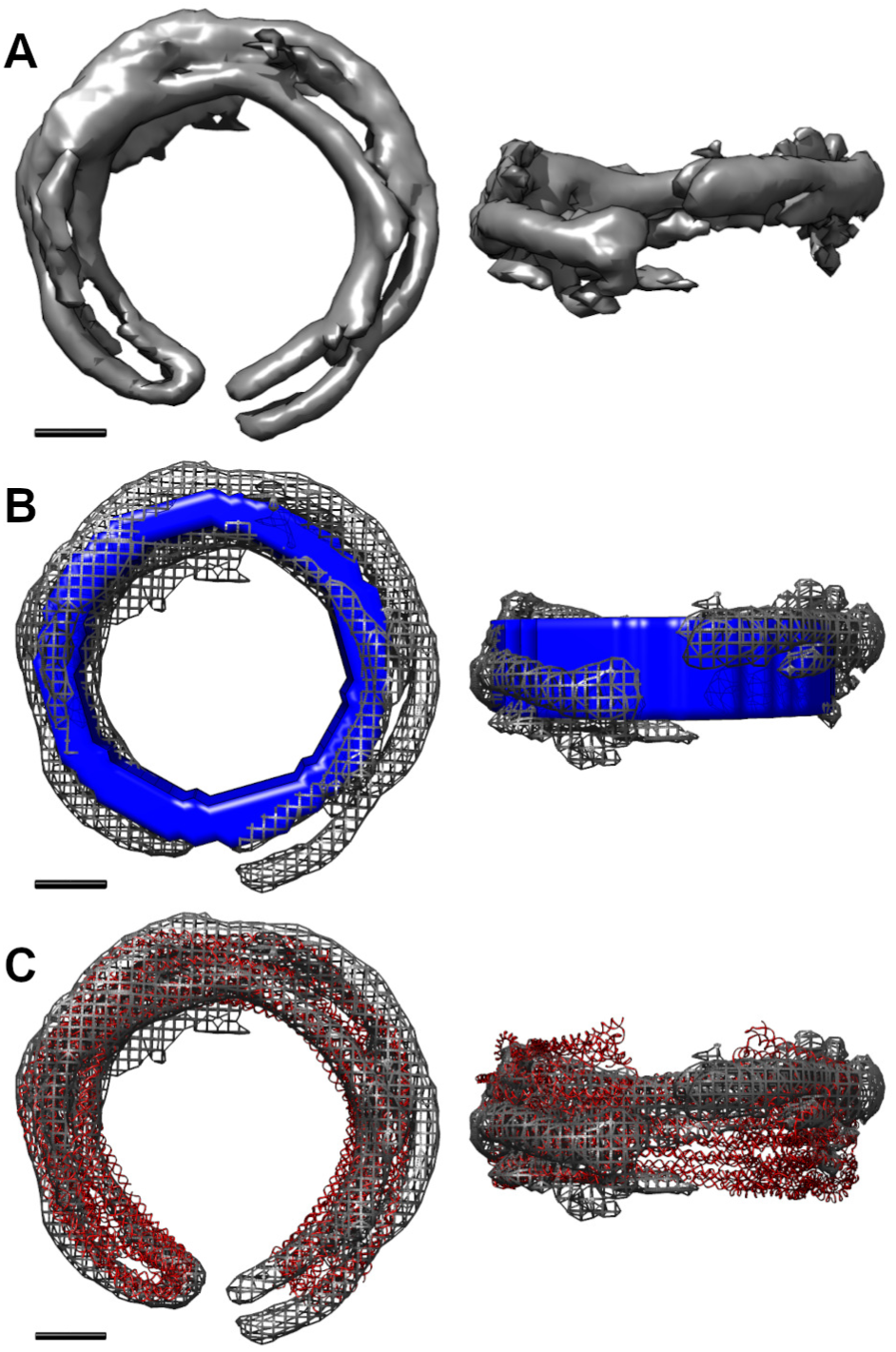
Experimental validation of the functionality of AutoMod with an example structure. (A) The reconstructed volume density map is shown as a solid grey surface. (B) The ideal target shape, without a gap, is illustrated with a solid blue surface. The reconstructed volume is displayed as a grey mesh, aligned with the target. (C) The predicted equilibrium conformation, obtained as the mean of *N* = 10 MD simulations, is shown as a red ribbon model, aligned with the mesh representation of the reconstructed volume. Scale bars are 10 nm.

### Coarse-Grained Molecular Dynamics Simulations

The oxDNA simulation engine [54, 55] with the DNA2 force field [56] was used for all coarse-grained molecular dynamics (MD) simulations. The equilibrium conformations for all versions of the split ring benchmark structure (Figures 2 and 5) were predicted by the following simulation protocol. First, preliminary relaxation was performed with 2000 steps of gradient descent with a timestep of 3.03 fs (0.001 in simulation units). Next, an MD simulation of 50 million steps was used to allow structural deformations, such as the bending of helices, to take place. A modified formulation for the backbone potential was used to ensure numerical stability during this step. Then, the modified formulation was gradually altered toward and finally replaced by the true backbone potential (Supplementary Note 5) over a total of 30 million steps. Lastly, a simulation of 100 million steps was performed, and the observed conformation was saved after every one million steps. An estimate of the equilibrium conformation was obtained by taking the mean over the 100 saved conformations. A timestep of 15.15 fs, a temperature of 303 K, a Newtonian step of 103, a John thermostat, a salt concentration of 0.5 M, and average sequence parameters were used in all MD simulations. The oxDNA analysis tools Python package [57] was used to calculate mean conformations.

The equilibrium conformations in Figures 3 and 4 were estimated with an otherwise identical simulation protocol, but the length of the first MD simulation was increased to 150 million steps to allow all major deformations to take place before subsequent simulations. Conversions from the caDNAno file format to the oxDNA format were done with the TacoxDNA [58] converter integrated into the oxView tool [59].

Initial gradient descent was performed on a CPU (13th Gen Intel Core i5-13400 × 16) on a Linux workstation (Linux 6.17.0-1032-oem, 64-bit, Ubuntu 24.04.4 LTS). OxDNA version 3.7.0 was compiled with GCC 11.2.0 and CMake 3.28.3.

An ensemble of either *N* = 10 (Figures 3-5, and Supplementary Figures S8-9), or *N* = 5 (Supplementary Figures S1-7), simulation repeats with different initializations of the random number generator was performed for each structure. Random seeds 1-10, or 1-5, were used for the repeats, respectively.

MD simulations following initial gradient descent were performed on GPUs (NVIDIA H200, H100, A100, V100) using oxDNA version 3.7.0, compiled with GCC 13.3.0, CMake 3.30.5, and CUDA 12.6.2. The Aalto University high-performance computing cluster, Triton, was used to parallelize the MD simulations of each ensemble.

### Quantification of Design Accuracy

The mean conformation for each structure, computed from MD simulation results, was further processed by finding the centroid of all base pairs. Each helix was then represented as the sequence of centroids for the base pairs belonging to the helix in question. Since helices in bent structures can have unequal total numbers of base pairs, the lengths of centroid sequences were equalized by downsampling all sequences to match the length of the shortest sequence. Then, centroid locations were averaged over all helices to obtain one set of points representing the full conformation. This roughly corresponds to taking the location of each helix in a series of cross section planes that have been placed uniformly over the full length of the structure and finding the centre of mass in each cross section plane.

Similarly, the target shape for each structure was represented by a point set of equal size. First, the target locations for the helical axes were represented by splines, and a sequence of points was sampled from the corresponding spline for each helix. Again, the final point set was obtained by averaging over the helices.

All point sets were centred on the origin by subtracting the centroid. Point sets representing simulation results were aligned to point sets representing the corresponding target shapes by minimizing the Root Mean Square Deviation (RMSD) with the SciPy library implementation [62] of the Kabsch algorithm. The accuracy of CAD software was then measured by the final RMSD between aligned point sets. A distance threshold (*d*_lim_) was used to categorize every pair of aligned points as either being close enough to each other (*d*(*a, b*) *≤ d*_lim_ =*⇒* (*a, b*) *∈ A*) or being too far away from each other (*d*(*a, b*) *> d*_lim_ =*⇒* (*a, b*) *∈ B*). Here, *d*(*a, b*) denotes the Euclidean distance between points *a ∈* ℝ^3^ and *b ∈* ℝ^3^. Accuracy as a percentage was then calculated from the set sizes as; Accuracy 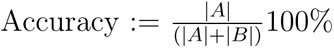. (See Sup-plementary Figure S10 for a schematic workflow that illustrates the process described above.)

The amount of run-to-run variation in MD simulations was quantified with the distribution of RMSD values between the point set representations from different MD runs within the same ensemble. Statistical testing for differences in design accuracy across different software and methods was conducted using a SciPy library implementation [62] of Welch’s t-test (Student’s t-test for two independent samples with unequal variances). The alternative hypothesis that the AutoMod designs have a lower mean RMSD than the other designs under consideration was tested against the null hypothesis that AutoMod designs have an equal or larger mean RMSD. (See Supplementary Figures S8-9 for additional accuracy comparisons.)

### Synthesis and Purification of DNA Origami Structures

A reaction mixture containing 10 nM of M13mp18 scaffold strands (Tilibit Nanosystems, Germany, Product No. M1-11), 100 nM of each staple strand (Integrated DNA Technologies, Belgium, Custom Oligos), 1X of TE (10 mM of Tris and 1 mM of EDTA) buffer (Fisher BioReagents, UK, Product No. BP1338-1), as well as 10 mM of MgCl_2_ (Sigma-Aldrich, Germany, Product No. M9272) was prepared by mixing all constituents. (See Supplementary Data 1 for staple strand sequences.) Self-assembly was performed by first raising the temperature of the reaction mixture to 85 *^◦^*C for 15 minutes and then slowly decreasing the temperature from 65 *^◦^*C to 40 *^◦^*C with a rate of one degree Celsius per hour in a PCR machine (Thermo Fisher Scientific, Applied Biosystems, ProFlex PCR System).

Initial purification was done with gel electrophoresis, for which an agarose gel containing one mass percent of agarose (Thermo Fisher Scientific, USA, Invitrogen UltraPure Agarose, Product No. 16500100), 0.5X of TBE (50 mM of Tris, 45 mM of Boric acid, and 0.5 mM of EDTA) buffer (Thermo Fisher Scientific, USA, Invitrogen UltraPure 10X TBE Buffer, Product No. 15581044), 13 mM of MgCl_2_, and 1X of SYBR Safe dye (Thermo Fisher Scientific, USA, Invitrogen SYBR Safe DNA Gel Stain, Product No. S33102) was prepared by dissolving the agarose powder in a heated buffer solution, adding other components, and letting the mixture solidify while cooling down. Unpurified reaction mixtures were then electrophoresed, while submerged in a running buffer containing 0.5X of TBE and 11 mM of MgCl_2_, with a bias voltage of 80 V 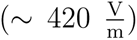 for approximately three hours in a chamber immersed in an ice-water bath. Leading monomer bands were excised from the gel under ultraviolet illumination, sliced with a razor blade, crushed with a glass slide, and passed through spin column filters (Bio-Rad Laboratories, USA, Product No. 7326165) by centrifuging for 5 min at 3000 RCF at room temperature.

For yield estimation, the leading monomer band was imaged (Bio-Rad Laboratories, USA, Gel Doc XR+ System) with ultraviolet light (*λ* = 302 nm). The background corrected, integrated intensity over the leading monomer band was compared to the background corrected, integrated intensity over the full lane to obtain a yield estimate (Supplementary Figure S11) of 71%.

Subsequent concentration adjustment was done with centrifugal filtering through 100 kDa molecular weight cutoff filters (Sigma-Aldrich, Germany, Amicon Ultra, Product No. UFC510096). The filters were first wetted by filtering 0.5 mL of buffer solution (0.5X of TBE and 11 mM of MgCl_2_) at 14000 RCF for 3 min. The gel purified origami samples were concentrated to approximately 10 nM by filtering at 3000 RCF using 10 min rounds and pooling the concentrated origami solution until the final concentration was reached. Concentrations were measured with ultraviolet-visible spectroscopy (Eppendorf, Germany, BioSpectrometer) by subtracting the background from the absorption at *λ* = 260 nm and applying the Beer-Lambert law, with 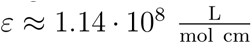 as the molar extinction coefficient of DO structures.

### Cryogenic Transmission Electron Microscopy

Sample aliquots of 5 *µ*L were pipetted onto holey carbon coated copper grids with a hole diameter of 1.2 *µ*m, hole spacing of 1.3 *µ*m, mesh size of 200 (grid squares per inch), and an ultrathin carbon support film (Quantifoil, R 1.2/1.3, UT, 200 Mesh, Cu) inside the temperature and humidity controlled chamber of the vitrobot (Leica Microsystems, Germany, Leica EM GP2). Temperature was set to 22 *^◦^*C and relative humidity to 90%. Grids were prepared by glow discharging with a plasma cleaner (Fischione Instruments, USA, NanoClean Model 1070) using 30% power (*∼* 11 *−* 12 W of forward power) and a gas mixture of oxygen and hydrogen (*vol*%(*O*_2_) : *vol*%(*H*_2_) = 75 : 25) with a gas flow rate of 20 sccm for 30 s. Sample aliquots were incubated for 90 s before backblotting the grids with filter paper. A blot time of 3 s was used. Blotted grids were immediately plunged into liquid ethane, cooled down by liquid nitrogen.

Vitrified samples were imaged with a cryo electron microscope (JEOL, Japan, JEM-3200FSC) equipped with a direct electron detector (Direct Electron, USA, Apollo) and operated at an acceleration voltage of 300 kV. A nominal magnification of 25000, resulting in a pixel size of 1.18 Å, was used for data collection. In total, 1116 movies were collected from one sample by recording 61 frames during a total exposure of 1 s at each imaging location. An objective aperture with a diameter of 50 *µ*m was used in conjunction with the in-column energy filter, which was centred at the zero-loss peak with a slit width of 30 eV. Total dose per movie was approximately 1.5 e*/*Å^2^.

### Single-Particle Analysis

The collected movies were imported to CryoSPARC [63], where dose weighting and patch motion correction were performed before merging the frames of each movie to obtain accumulated frames. After patch Contrast Transfer Function (CTF) estimation, a total of 1193 particles were manually picked from 54 micrographs. Templates for automated particle picking were created by performing 2D classification with the manually picked particles. Following template picking, 157432 particles were extracted from the full set of micrographs. 2D classification was performed with all particles, after which 6070 picks were discarded. An initial consensus volume was created using ab-initio reconstruction and homogeneous refinement with all remaining particles. After inspecting the distribution of per-particle scale factors, a subset of 19443 particles was discarded due to low scale factors. For the remaining 131764 particles, ab-initio reconstruction and homogeneous refinement were performed again. Lastly, 3D flexible refinement was used to improve the resolution of the obtained consensus volume. (See Supplementary Figure S12 for further details on the data processing steps.) The resolution of the final volume density map was quantified using Gold-Standard Fourier Shell Correlation (GSFSC), which resulted in a resolution estimate of approximately 19 Å (Supplementary Figure S13).

## Results

### Comparison of Design Accuracy Across Methods and Software

Design accuracies for the benchmark structure across different design methods and software are displayed in Figure 2. It can be seen that the software which track the rotational phases of helices and preferentially place crossovers into positions with compatible phases output designs without large unwanted torsion out of the osculating plane. These software include ENSnano, DNAxiS, and AutoMod. Some torsion can be seen in the output of MagicDNA and in the manual design. This is as expected, since out-of-phase crossover positions are utilized in these designs.

Besides unwanted torsion, another notable feature in the results in Figure 2 is that many software output designs with a magnitude of curvature that is clearly below the target value. That is, instead of a split ring with the ends close to touching each other, many software output designs that resemble half rings. Presumably, this is because the density of the crossover pattern is not taken into account in the calculation of helix lengths, leading to an unaccounted loss of global curvature.

To further evaluate the design accuracy of AutoMod, a collection of structures with target shapes containing different magnitudes of curvature and utilizing a diverse variety of cross-sectional profiles, both in the honeycomb and the square lattice arrangements, were designed. For comparison, structures with the same target shapes were also designed with MagicDNA, which is, according to the authors current knowledge, the only previously published software capable of fully automated design of varied curvature, including automated crossover placement. A gallery of results can be seen in Figures 3-4. Supplementary Figures S8-9 contain additional results for more challenging target shapes.

In the comparisons included in Figures 3-4, AutoMod offers statistically significant decreases in the differences between the targets and the predicted equilibrium conformations of designs with all tested target shapes. The relative percentage changes in the mean RMSD values are approximately *−*53% (honeycomb lattice) and *−*59% (square lattice) for the target shapes in Figure 3, and approximately *−*62% (honeycomb lattice) and *−*80% (square lattice) for the target shapes in Figure 4.

The main factors presumably contributing to the increase in accuracy are the spatially more finely discretized solution for length difference placement, i.e., targeted modification placement is used instead of dividing the structure into large linear segments, the inclusion of modifications in the tracking of the rotational phases of the helices, and the introduction of crossover density dependent modification calculation. The design method implemented by AutoMod appears to generalize quite well across different shapes and cross sections.

However, the design of structures containing extremely high curvature, with a radius of curvature *R ≤* 10 nm, and rapid changes in curvature (Supplementary Figures S8-9) appears to still suffer from limited accuracy. Since only initial outputs, generated with the default settings are considered, accuracies might be further improved by structure-wise fine-tuning of model parameters and iterative manual refinement.

### Observed Conformation of the Example Structure

To experimentally confirm the high design accuracy of AutoMod, a structure similar to the split ring benchmark structure in Figure 2 was designed with a slightly increased radius (*R* = 21.8 nm) to closely match the size of the M13mp18 scaffold strand. After synthesis and purification, the equilibrium conformation was resolved with single-particle reconstruction using cryoTEM data. The results (Figure 5) suggest a high similarity between the ideal target shape, the obtained reconstruction, and the mean result of *N* = 10 different MD simulation runs, thus supporting the previous computational findings.

The final resolution of approximately 19 Å appears to be sufficient for resolving the gap between the ends of the split ring, whose narrowness demonstrates that the magnitude of global curvature displayed by the structure is close to the target value.

## Discussion

As demonstrated in this work, the main advantage of the design approach implemented by AutoMod is increased design accuracy. High design accuracy is of great importance because, in most applications, the ability of DO structures to fulfill their desired functions depends on sufficient similarity between their equilibrium conformations and the intended target shape. Especially the equilibrium conformations of strongly curved structures whose helices span non-closed routes have been prohibitively challenging to design, limiting the application potential of the DO technique.

Besides the main advantage of increased accuracy, the limitations of the here presented approach should also be highlighted. They include the potential irregularity of generated crossover patterns, the sensitivity of the solutions to model parameters, and the fact that the dependence of the effectiveness of modifications on crossover spacing is only acknowledged at a high level by controlling the average crossover density of each helix, instead of individual control over the length of each helical segment in the structure.

The potential irregularity of crossover patterns follows from the preferential placement of crossovers to positions where the sugar-phosphate backbones of the strands that switch helices are spatially as close to each other as possible. This minimizes the magnitudes of the twist deformations that the helices have to undergo for the structure to successfully realize all designed crossovers. However, for some target shapes, the preferential selection of well-aligned crossover positions coupled with the control of crossover density only at the level of helix-wise averages can result in crossover patterns where the distances between consecutive crossovers, and the local crossover density, vary significantly within a helix. For applications prioritizing the internal homogeneity of structures, this is unfavourable.

Another limitation of the presented approach is its sensitivity to the parameter values acting as global constants. Accumulation of error can occur if incorrect parameter values are used, rendering high-accuracy outputs conditional on correct choices for parameter values. Thus, even though the wide applicability of the default settings is demonstrated here, it should be emphasized that in some cases the end user may have to adjust the settings to obtain the most accurate output. Similarly, if DNA structures are designed for applications that take place in solution conditions notably different from the ones described here, this should be taken into account in the design phase. We strongly encourage users to utilize computational methods, such as coarse-grained molecular dynamics simulations or finite element analysis, to evaluate the equilibrium conformations of structures designed with AutoMod prior to any experimental work.

Here, a straightforward approach is proposed for addressing the dependence of the efficiency of modifications on the crossover spacing. In summary, one model parameter (*δ*) is dedicated to capturing the effective length change caused by the placement of one modification, and another parameter (*ρ*) to describing the crossover spacing. Both parameters are predetermined constants, and high design accuracy can be achieved by finding such a pair of values that any decrease in the effectiveness of modifications due to a limited crossover density is intentionally compensated for by an increase in the total amount of modifications.

This is a fairly crude approach that could be replaced by a more sophisticated solution method by allowing the value of *δ* to vary between different locations, instead of treating it as a global constant. However, if modifications were assigned different effective length change values depending on the positions of the nearby crossovers, a more demanding solution procedure would likely be required since the modification pattern would have to be solved iteratively in conjunction with the crossover pattern. It should also be mentioned that in this work, the outputs of the CAD software were minimally altered for the sake of an objective accuracy comparison. Thus, it may be possible to achieve higher design accuracies than the ones demonstrated here by iterative refinement of designs based on feedback from molecular dynamics simulations or finite element analysis.

Lastly, some limits for the space of feasible designs are set by the properties of DNA. Assuming that the induced bend and other deformations are elastic, the total potential energy stored in these deformations must not exceed the amount of work available via the energetically favourable hybridization reactions between complementary sequences. In addition, the number of well-aligned crossover positions where participating helices have suitable rotational phases is limited, and set by the selected target shape along with the selected cross-sectional profile.

In conclusion, AutoMod offers a refined method for the computer-aided design of curved DNA nanostructures. AutoMod aims to ease the design of continuously curved structures with challenging target shapes by producing initial outputs with high accuracy. We anticipate that AutoMod may facilitate the actualization of structures that have previously remained unattainable, thus enabling future advances in the field of structural DNA nanotechnology.

## Data Availability

We distribute AutoMod as a Python package registered to the Python Package Index (PyPI, https://pypi.org/) under the package name *automod*. Version 0.1.4 was used for design generation. For reproducibility, snapshots of the internal state of the program prior to each design generation are provided in the supporting files. The source code is also available in several public repositories, including Zenodo (https://doi.org/10.5281/zenodo.22751375), Figshare (https://doi.org/10.6084/m9.figshare.30882791), and GitHub (https://github.com/AutoMod-DNA/automod.git). The source code is released under an MIT licence and no costs to the end user are associated with its acquisition or use. CaDNAno files for all designs, oxDNA files for all mean conformations, internal state files for AutoMod, and all code for processing data and creating figures are publicly available on GitHub (https://github.com/AutoMod-DNA/automod__supporting_files). The reconstructed volume density map is publicly available in the Electron Microscopy Data Bank (EMDB) with the identifier EMD-100132.

## Supplementary Information

Supplementary Information is provided as a separate file.

## Author Contributions Statement

E.H., A.K.N., and An.K. conceived the design method and the experiments. E.H., T.L., and Ab.K. developed the software. A.K.N. provided guidance for cryoTEM and single-particle reconstruction. E.H. conducted the experiments, analysed the results, and wrote the manuscript. All authors reviewed and edited the manuscript. An.K. supervised the project.

## Acknowledgments

The authors acknowledge the provision of facilities and technical support by Aalto University at OtaNano - Nanomicroscopy Center (Aalto-NMC), and the computational resources provided by the Aalto Science-IT project. We thank Professor Luc St-Pierre for discussions on developing the design method.

