## Supplementary Information for "AutoMod: A Software Tool for High-Accuracy Design of Curved DNA Origami Nanostructures"

This PDF file includes:

**Supplementary Figure S1**
Dependence of global curvature on crossover density
**Supplementary Figures S2-3**
Calibration of default parameters
**Supplementary Figure S4**
Accuracy comparison between extrude and sweep methods
**Supplementary Figure S5**
Accuracy comparison for manually altered edge gradients
**Supplementary Figure S6**
Conformational variation due to staple nick placement
**Supplementary Figure S7**
Conformational variation due to randomness in the design process
**Supplementary Figures S8-9**
Accuracy comparisons for additional example structures
**Supplementary Figure S10**
Quantification of design accuracy
**Supplementary Figure S11**
Yield estimation
**Supplementary Figure S12**
Detailed single-particle reconstruction process
**Supplementary Figure S13**
Gold-standard Fourier shell correlation of the final reconstruction
**Supplementary Equations 1**
Discretization of arc length differences
**Supplementary Equations 2**
Manual modification placement
**Supplementary Equations 3**
Parametric equations and sample point sequences for all target curves
**Supplementary Note 1**
Crossover placement
**Supplementary Note 2**
DNAxiS settings
**Supplementary Note 3**
AutoMod default settings
**Supplementary Note 4**
Autobreak settings
**Supplementary Note 5**
Full oxDNA simulation protocol
**Supplementary Data 1**
Staple strand sequences

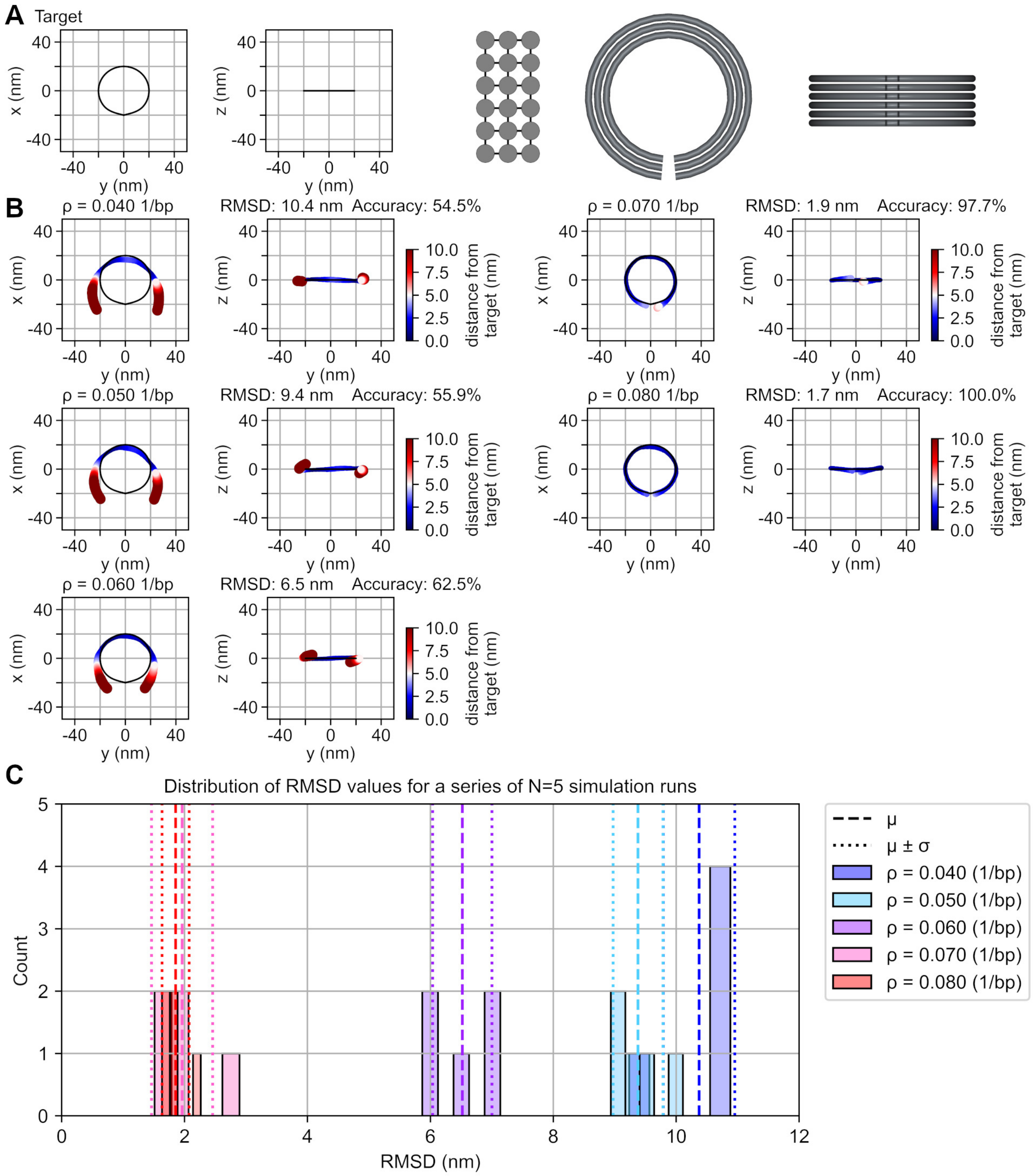


**Supplementary Figure S1:** Demonstration of the dependence of global curvature on crossover density. **(A)** The target shape. On the left, the point set representation for the target shape is shown. The cross-sectional profile is shown in the middle, with black lines connecting neighboring helices, in between which crossovers can take place. On the right, a drawing of the target shape with DNA helices as gray, homogeneous beams can be seen. **(B)** Accuracy evaluations for otherwise identical designs but with the target value for the average crossover density of each helix set to $0.040\frac{1}{\text{bp}}$, $0.050\frac{1}{\text{bp}}$, $0.060\frac{1}{\text{bp}}$, $0.070\frac{1}{\text{bp}}$, or $0.080\frac{1}{\text{bp}}$. The value of the effective length change induced by a modification was set to $\delta = 0.22$ nm for all designs. As crossover density is increased while holding $\delta$ constant, an increase in global curvature can be observed. Means over $N = 5$ simulation run repeats are shown. **(C)** Distribution of RMSD values over all $N = 5$ simulation repeats for each structure. Five bins of equal width are distributed uniformly between the minimum and maximum value of each distribution.

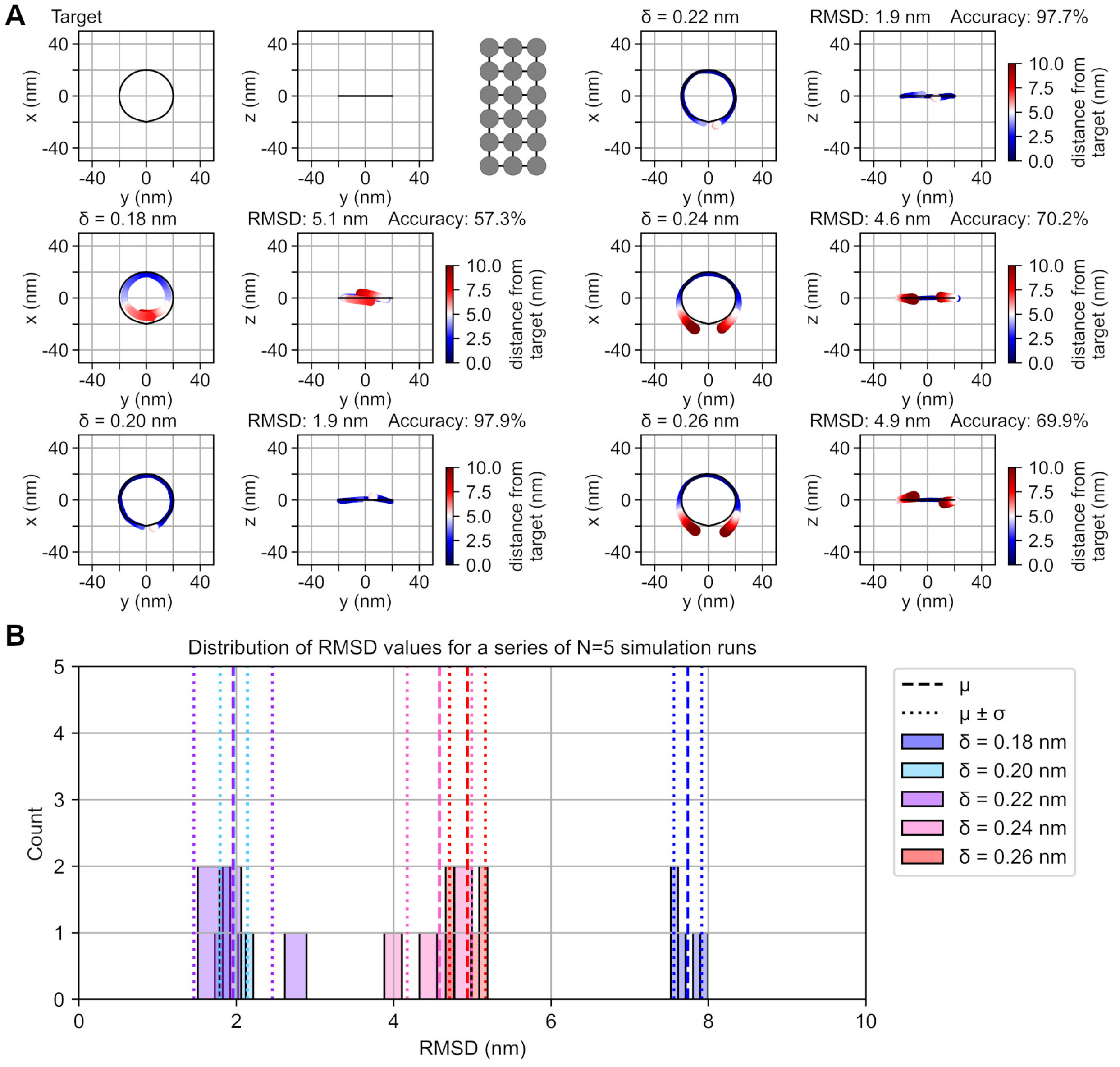


**Supplementary Figure S2:** Calibration of parameters using a square lattice cross section. One possible approach to selecting parameter values is depicted here and proceeds as follows: First, some obtainable target value for crossover density is chosen. Here, $\rho = 0.070\frac{1}{\mathrm{bp}}$ is used. Then, the value of the effective length change induced by a modification ($\delta$) is tuned to yield maximal accuracy. Here, a calibration series containing the test values 0.18 nm, 0.20 nm, 0.22 nm, 0.24 nm, and 0.26 nm are shown. Alternatively, $\delta$ could be held constant while sweeping over a series of values for crossover density. **(A)** The target shape. **(B)** Accuracy evaluations for structures in the calibration series. Means over $N = 5$ simulation run repeats are shown. **(C)** Distribution of RMSD values over all $N = 5$ simulation repeats for each structure. Five bins of equal width are distributed uniformly between the minimum and maximum value of each distribution.

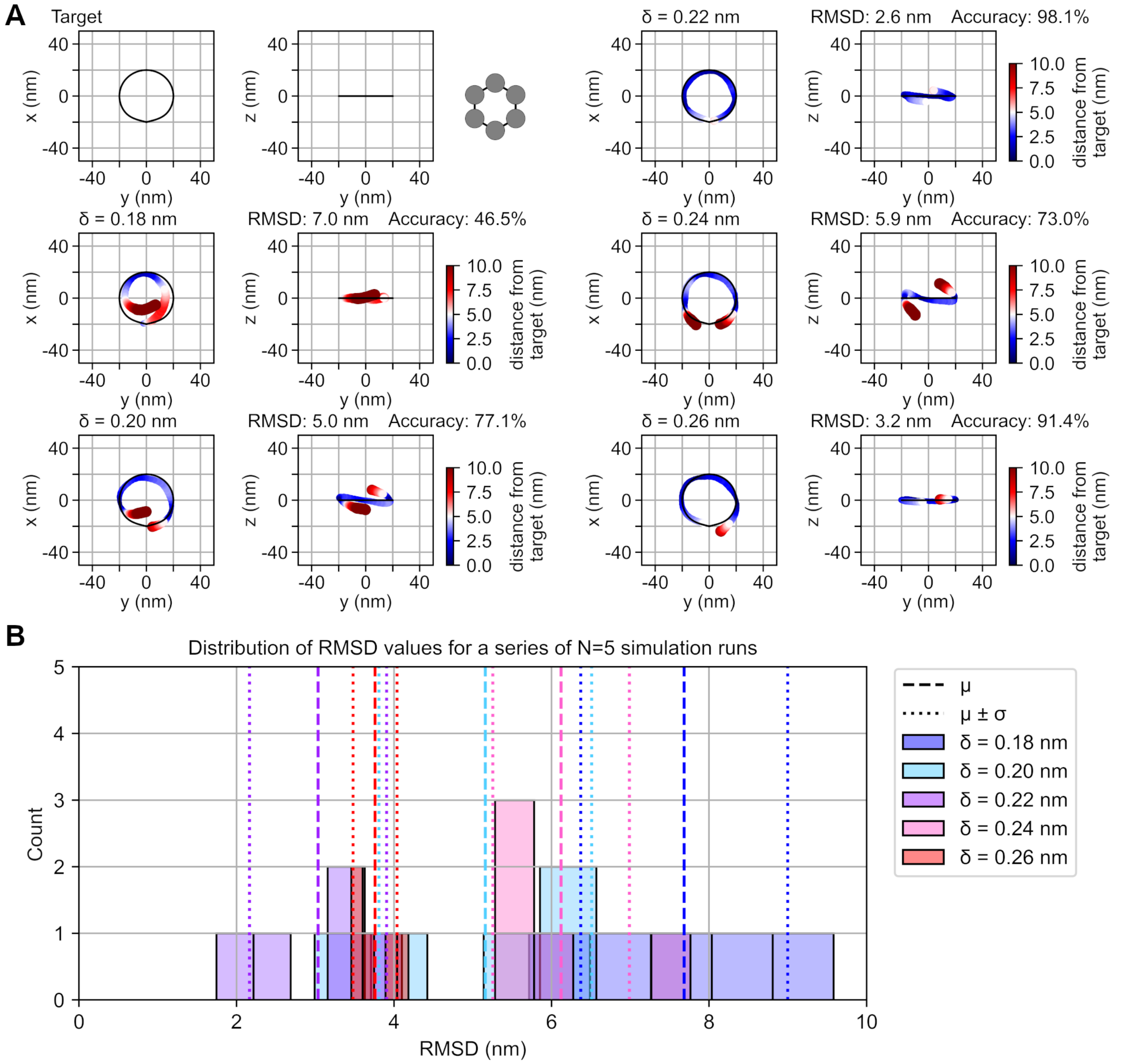


**Supplementary Figure S3:** Calibration of parameters using a honeycomb lattice cross section. One possible approach to selecting parameter values is depicted here and proceeds as follows: First, some obtainable target value for crossover density is chosen. Here, $\rho = 0.070 \frac{1}{\text{bp}}$ is used. Then, the value of the effective length change induced by a modification ($\delta$) is tuned to yield maximal accuracy. Here, a calibration series containing the test values 0.18 nm, 0.20 nm, 0.22 nm, 0.24 nm, and 0.26 nm are shown. Alternatively, $\delta$ could be held constant while sweeping over a series of values for crossover density. **(A)** The target shape. **(B)** Accuracy evaluations for structures in the calibration series. Means over $N = 5$ simulation run repeats are shown. **(C)** Distribution of RMSD values over all $N = 5$ simulation repeats for each structure. Five bins of equal width are distributed uniformly between the minimum and maximum value of each distribution.

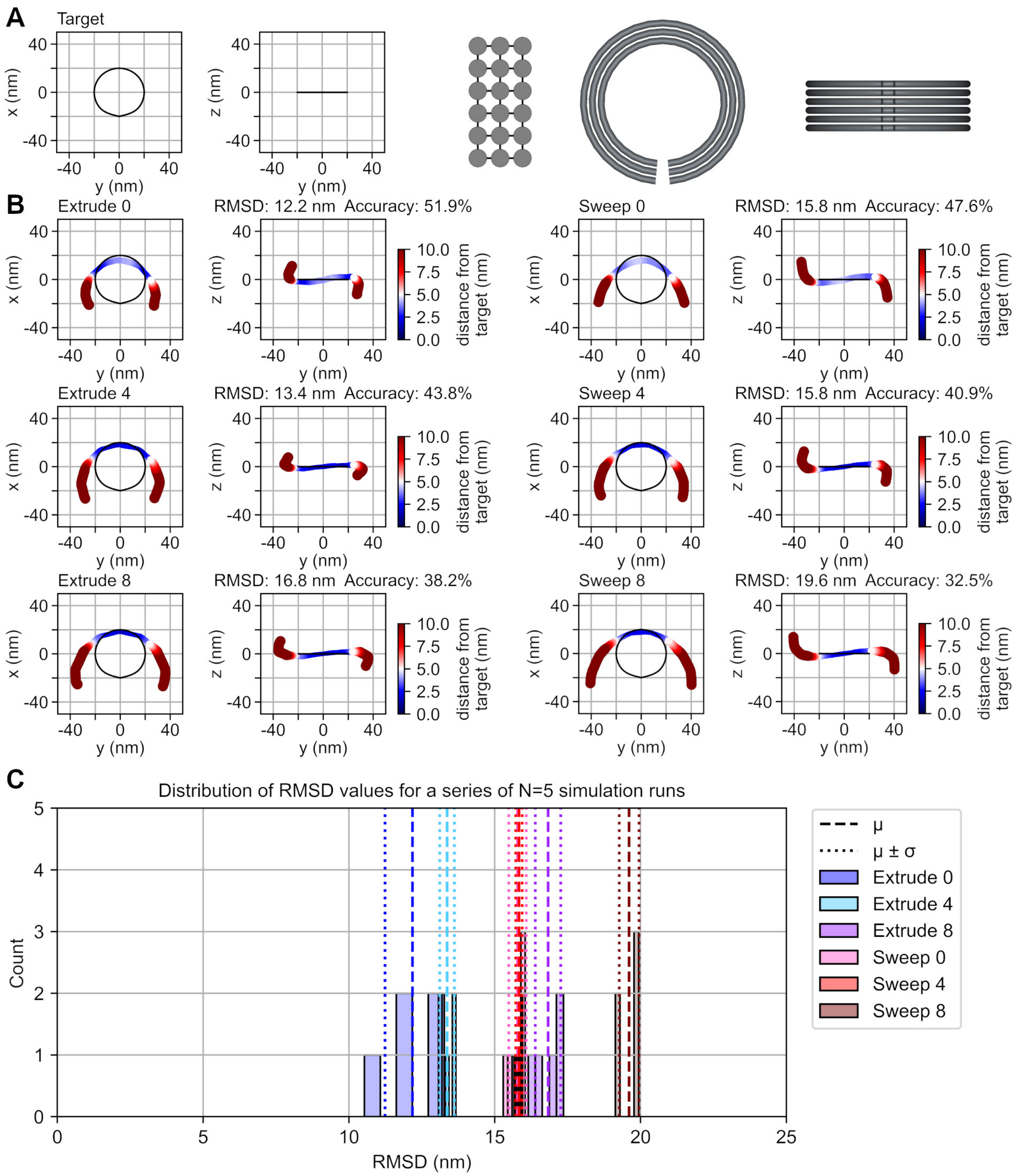


**Supplementary Figure S4:** Accuracy comparison for the two design methods for curved structures available in MagicDNA [1], i.e., the extrude and the sweep methods, with different numbers of spacer nucleotides in between linear segments, i.e., unpaired nucleotides in the portions of the scaffold strand that connect consecutive segments. **(A)** The target shape. **(B)** Accuracy comparisons with no spacer nucleotides in between segments, with 4 spacer nucleotides, and with 8 spacer nucleotides. A distance threshold value of; $d_{\mathrm{lim}} = 5.0$ nm, was used to compute all accuracy percentages. **(C)** Distribution of RMSD values over all $N = 5$ simulation repeats for each structure. Five bins of equal width are distributed uniformly between the minimum and maximum value of each distribution.

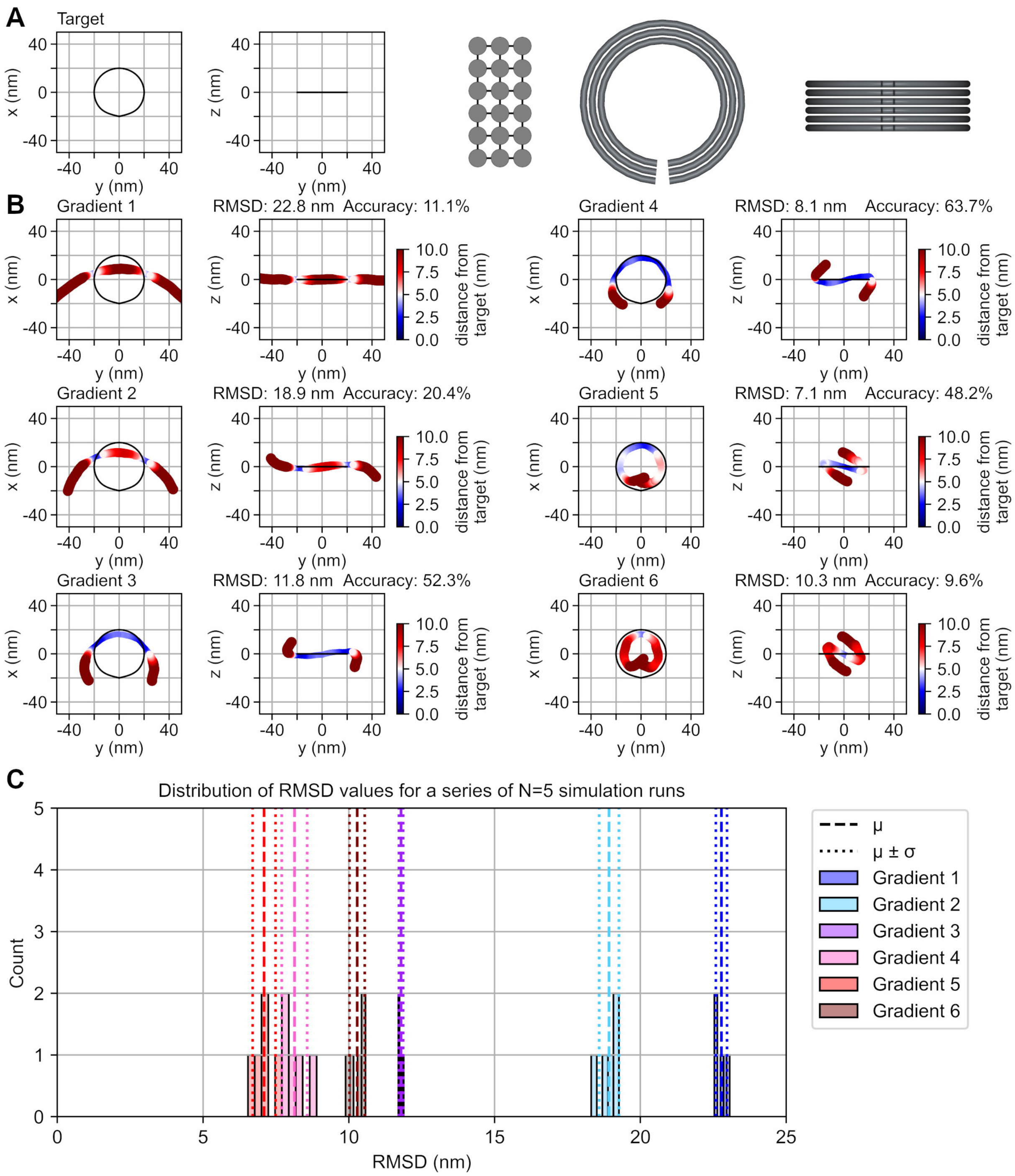


**Supplementary Figure S5:** Accuracy comparison between manually altered edge gradient magnitudes. **(A)** The target shape. **(B)** Accuracy evaluations for the designs where edge gradient magnitudes were manually set to 1 bp, 2 bp, 3 bp, 4 bp, 5 bp, and 6 bp. The default value without manual adjustment is $\sim$ 3 bp. A distance threshold value of; $d_{\lim} = 5.0$ nm, was used to compute all accuracy percentages. **(C)** Distribution of RMSD values over all $N = 5$ simulation repeats for each structure. Five bins of equal width are distributed uniformly between the minimum and maximum value of each distribution.

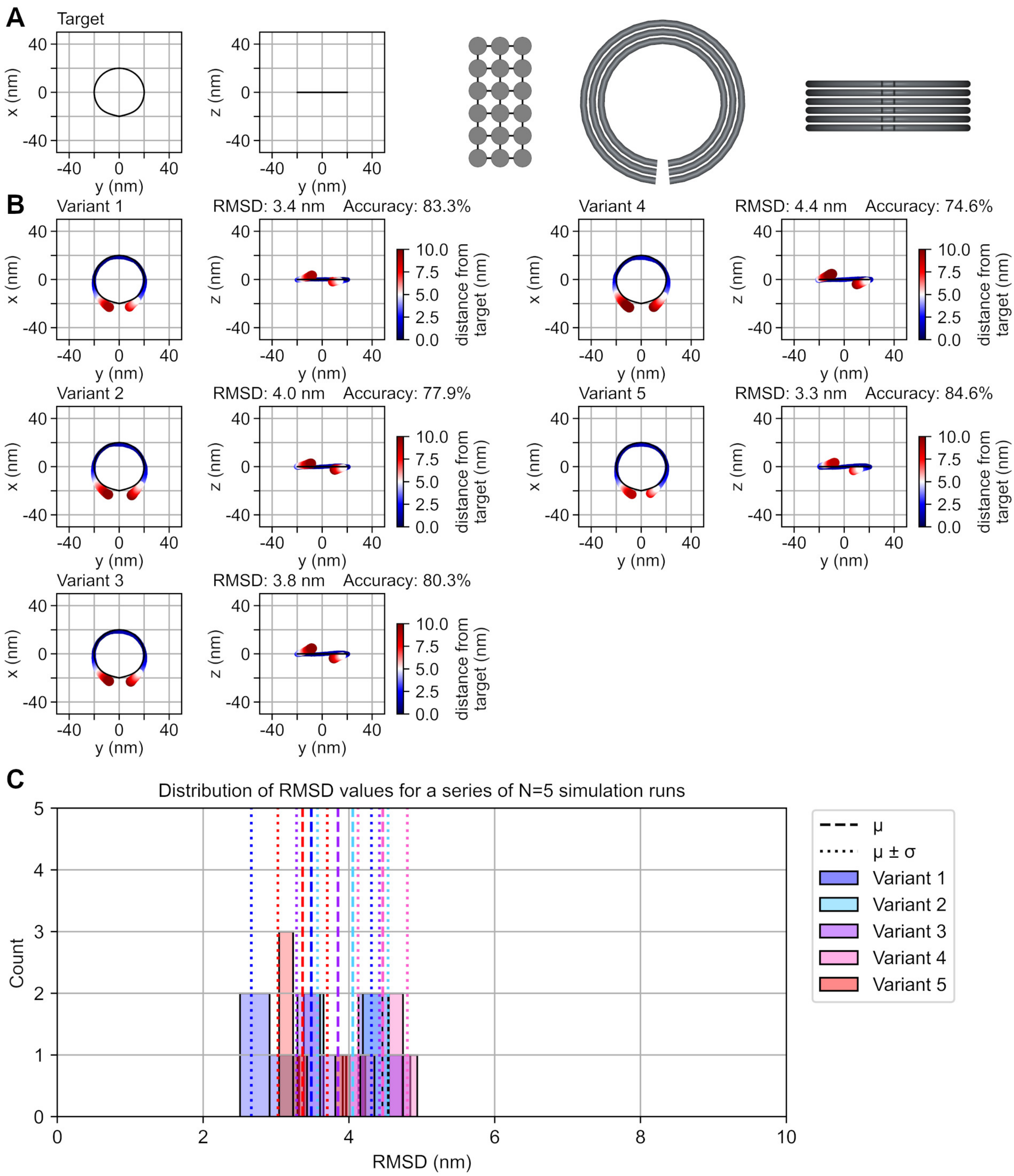


**Supplementary Figure S6:** Conformational variation due to staple nick placement. **(A)** The target shape. **(B)** Otherwise identical designs but with differently placed backbone nicks defining the start and end points of staple strands. Variants were generated with the Autobreak tool [2] by altering the random seed from 0 (Variant 1) to 4 (Variant 5). Minimum length of staples was set to 15 nucleotides. Otherwise, default settings were used. **(C)** Distribution of RMSD values over all $N = 5$ simulation repeats for each structure. Five bins of equal width are distributed uniformly between the minimum and maximum value of each distribution.

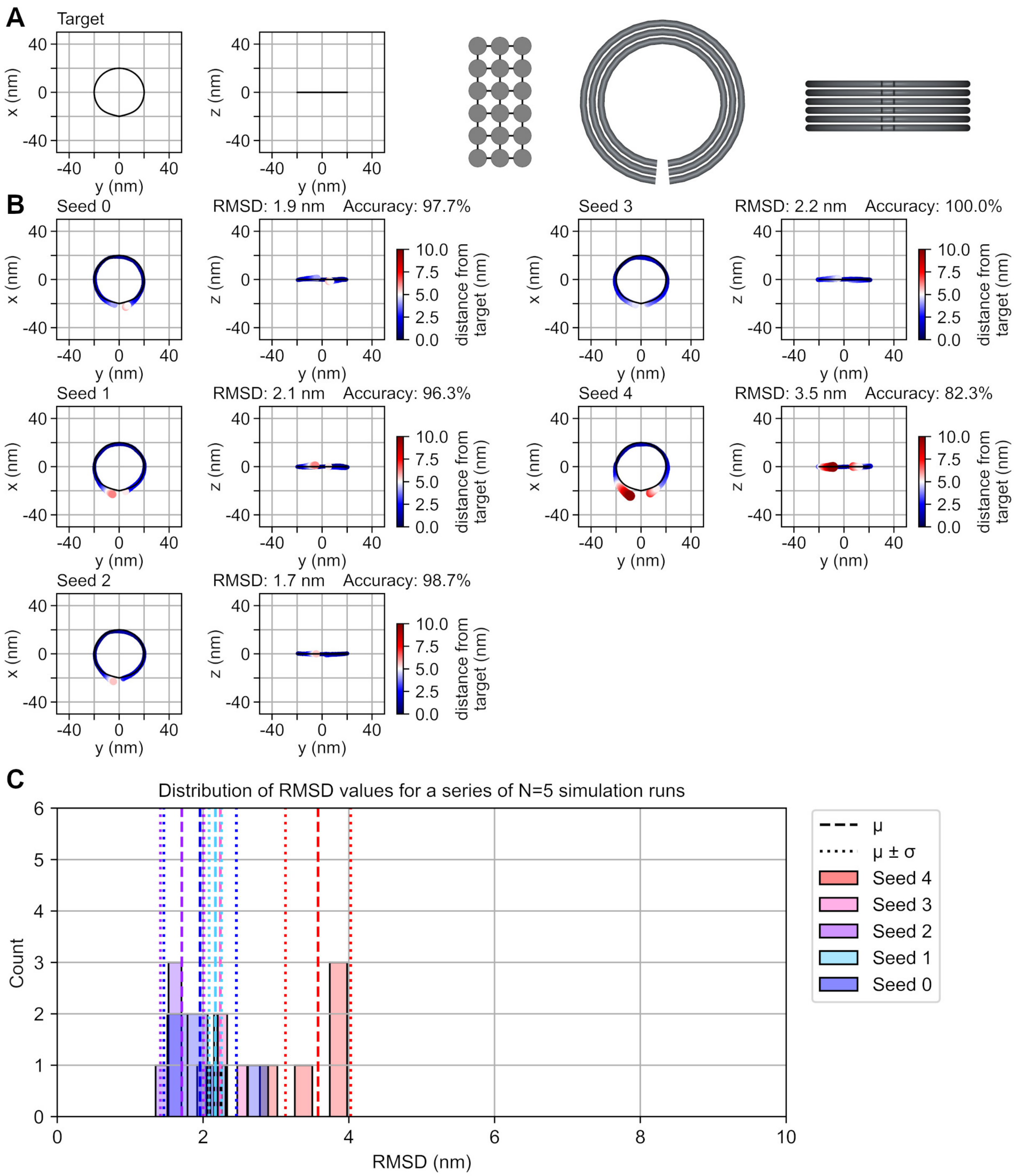


**Supplementary Figure S7:** Conformational variation due to randomness in the design process. **(A)** The target shape. **(B)** Otherwise identical designs but with different crossover patterns. The random seed used to initialize the random number generator in AutoMod was incremented from 0 to 4 to generate different crossover patterns. Otherwise, default settings were used. **(C)** Distribution of RMSD values over all $N = 5$ simulation repeats for each structure. Five bins of equal width are distributed uniformly between the minimum and maximum value of each distribution.

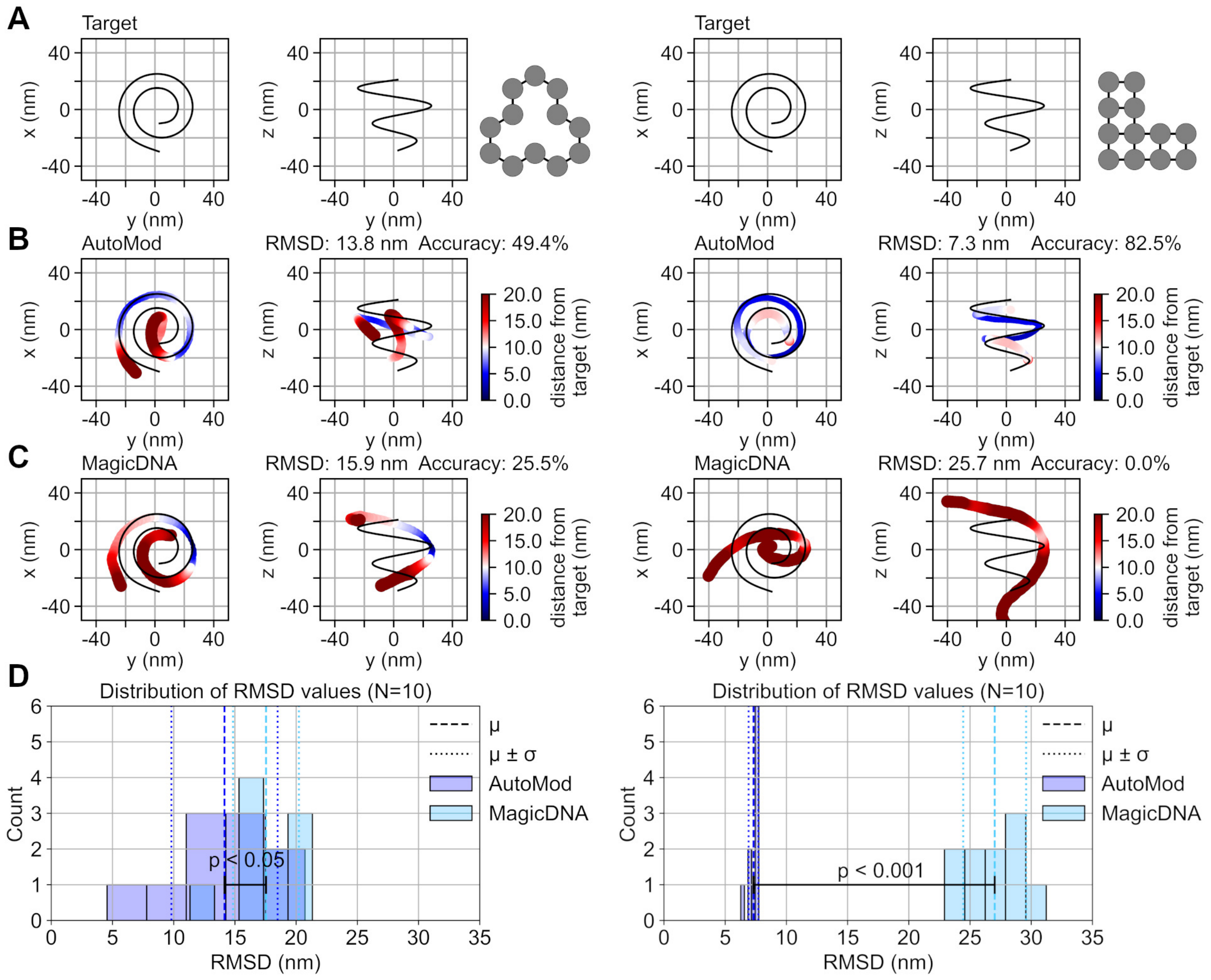


**Supplementary Figure S8:** Evaluation of design accuracy with a curved target shape and two different cross-sectional profiles adhering to the honeycomb and square lattices. A distance threshold value of; $d_{\mathrm{lim}} = 10.0$ nm, was used to compute all accuracies. **(A)** Point set representations of target shapes along with the cross-sectional profiles on the right. **(B)** Accuracy of AutoMod designs as given by the total mean over all simulation runs. **(C)** Accuracy of MagicDNA [1] designs as given by the total mean over all simulation runs. **(D)** Distributions of RMSD values for an ensemble of $N = 10$ different simulation runs for each structure. Five bins of equal width are distributed uniformly between the minimum and maximum value of each distribution. Welch's t-test confirmed the alternative hypothesis that the mean of the AutoMod distribution is less than the mean of the MagicDNA distribution, with p-values of approximately 0.03 (honeycomb lattice) and $7.3 \cdot 10^{-10}$ (square lattice).

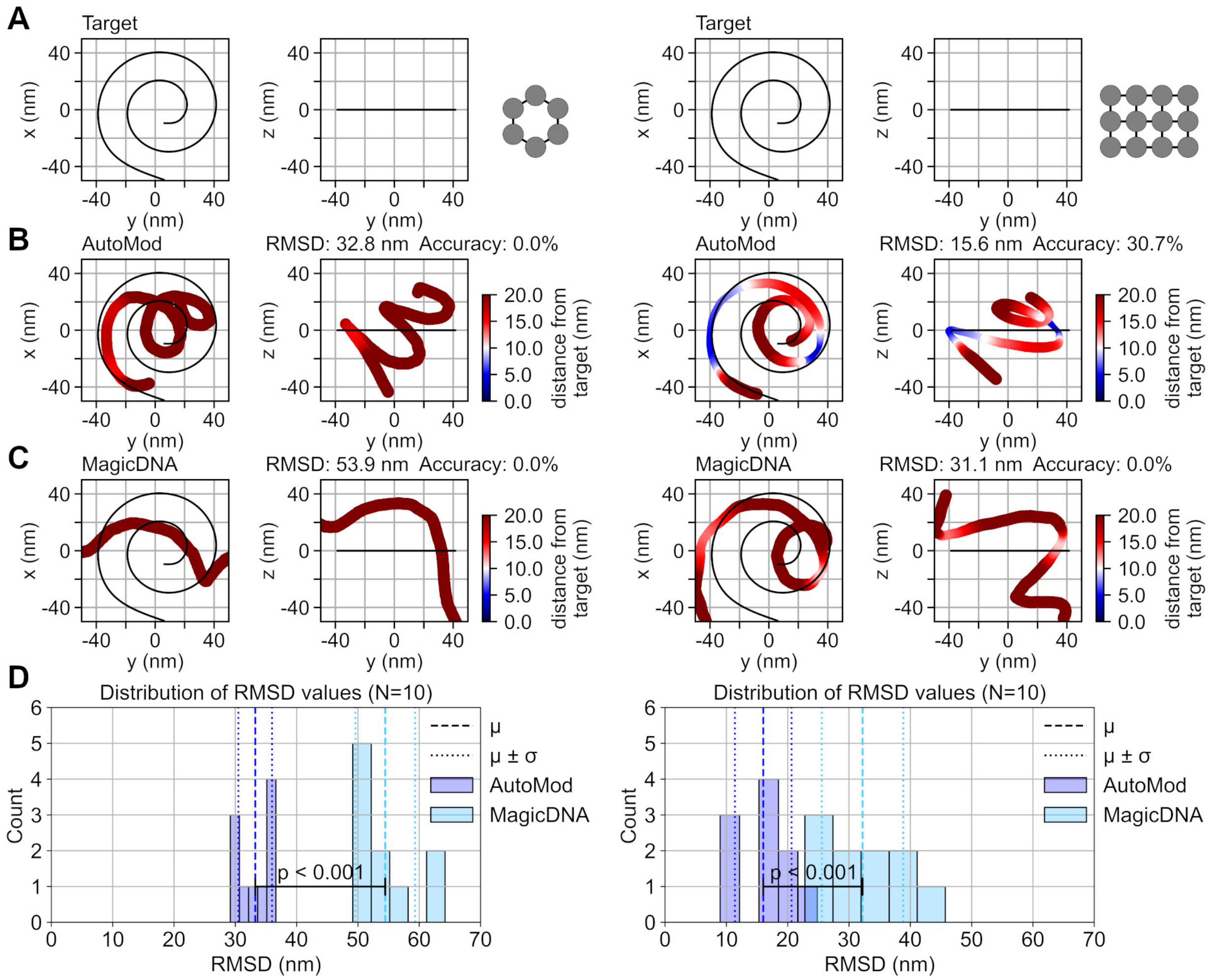


**Supplementary Figure S9:** Evaluation of design accuracy with a curved target shape and two different cross-sectional profiles adhering to the honeycomb and square lattices. A distance threshold value of; $d_{\text{lim}} = 10.0$ nm, was used to compute all accuracies. **(A)** Point set representations of target shapes along with the cross-sectional profiles on the right. **(B)** Accuracy of AutoMod designs as given by the total mean over all simulation runs. **(C)** Accuracy of MagicDNA [1] designs as given by the total mean over all simulation runs. **(D)** Distributions of RMSD values for an ensemble of $N = 10$ different simulation runs for each structure. Five bins of equal width are distributed uniformly between the minimum and maximum value of each distribution. Welch's t-test confirmed the alternative hypothesis that the mean of the AutoMod distribution is less than the mean of the MagicDNA distribution, with p-values of approximately $8.0 \cdot 10^{-9}$ (honeycomb lattice) and $9.2 \cdot 10^{-6}$ (square lattice).

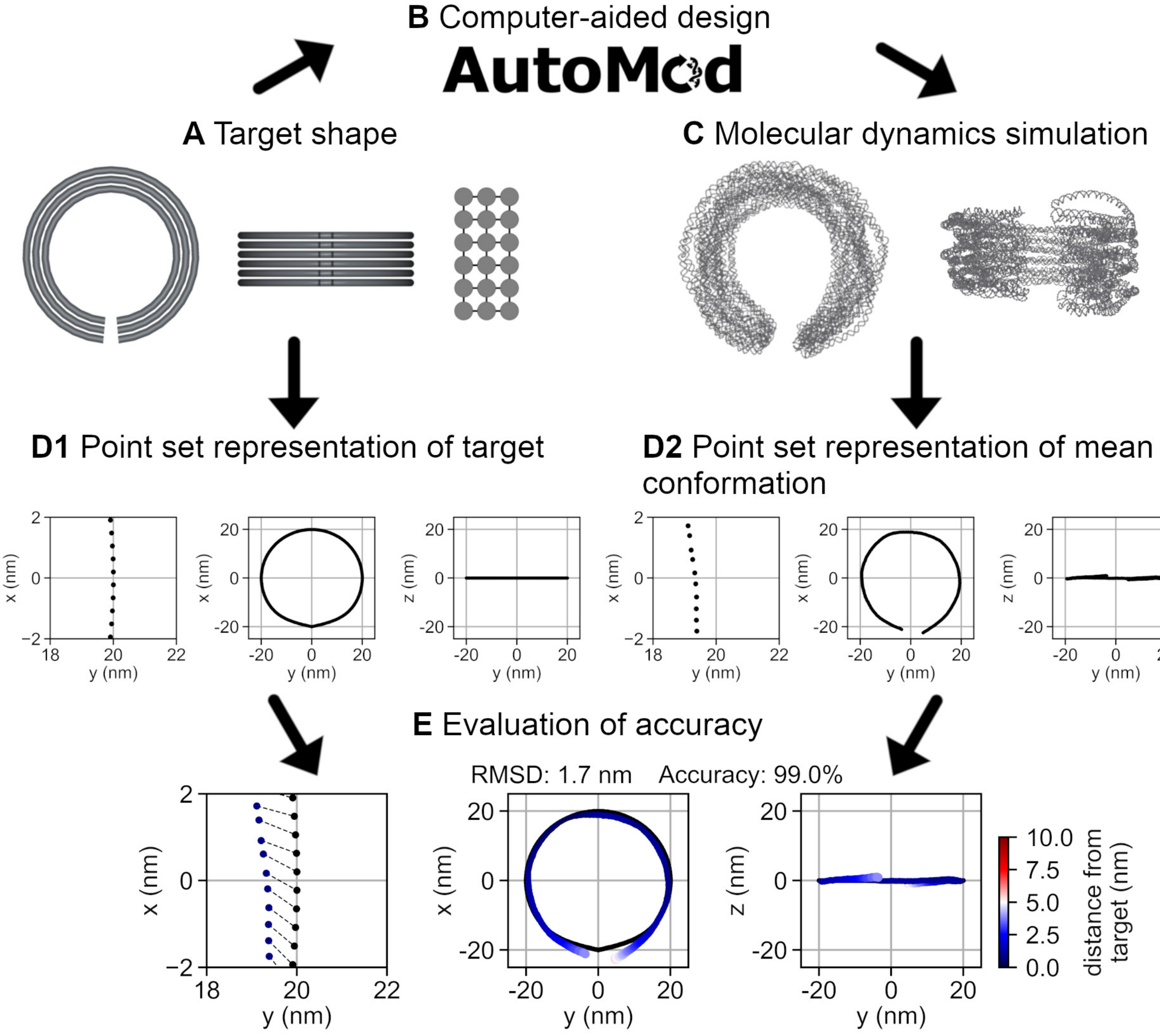


**Supplementary Figure S10:** Schematic workflow for the quantification and *in silico* evaluation of the design accury of different methods and CAD software. **(A)** First, a target shape is selected. **(B)** The target shape is inputted into CAD software that produces a strand arrangement aiming to realize the target shape as its equilibrium conformation. **(C)** Trajectories from coarse-grained molecular dynamics simulations are used to compute mean conformations by averaging over several consecutive conformations. Mean conformations are used as estimates of equilibrium conformations. **(D1)** Target shape is converted to a sequence of points by sampling the parametric curve that represents the route of the cross-sectional profile. **(D2)** Mean conformation is converted to a point sequence by finding the centroid of each basepair, downsampling centroid sequences to match the length of the shortest helix, and averaging over helices. **(E)** The Root Mean Squared Distance (RMSD) value for the best alignment between the simulation result and the target quantifies the accuracy of the design. The fraction of simulation result points that are at most the threshold distance away from their corresponding target points in the best alignment is used to calculate accuracy as a percentage. A threshold distance of $d_{\mathrm{lim}} = 5.0$ nm is used here.

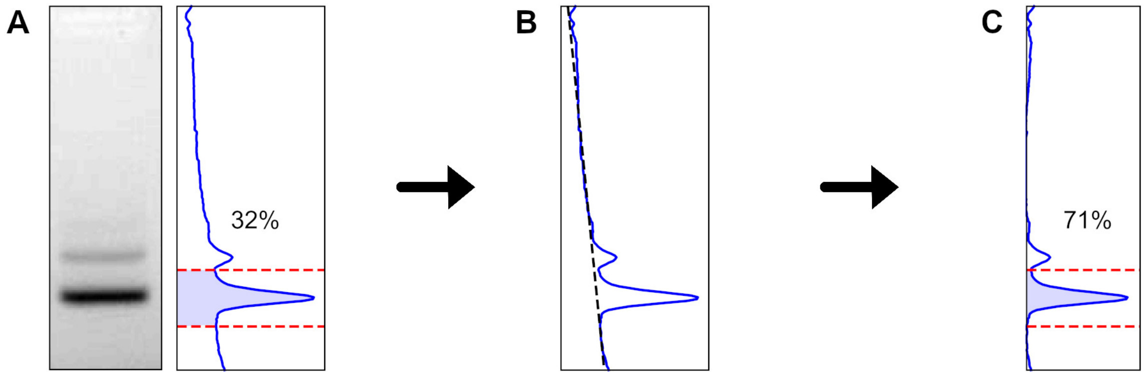


**Supplementary Figure S11:** Yield estimation from a gel image with linear background correction. **(A)** On the left, the lane containing the sample of interest is shown. A grayscale colormap from white to black, that shows maximum intensity as black, is used. On the right, the unprocessed intensity profile of the image displaying the lane is shown. The yield estimate of 32% is obtained by taking the ratio of the total intensity within the red dashed lines over the total intensity of the full image. **(B)** A linear fit to the background signal. **(C)** The final yield estimate of 71% is obtained by deducting the linear background from the intensity profile before taking the ratio of total intensities.

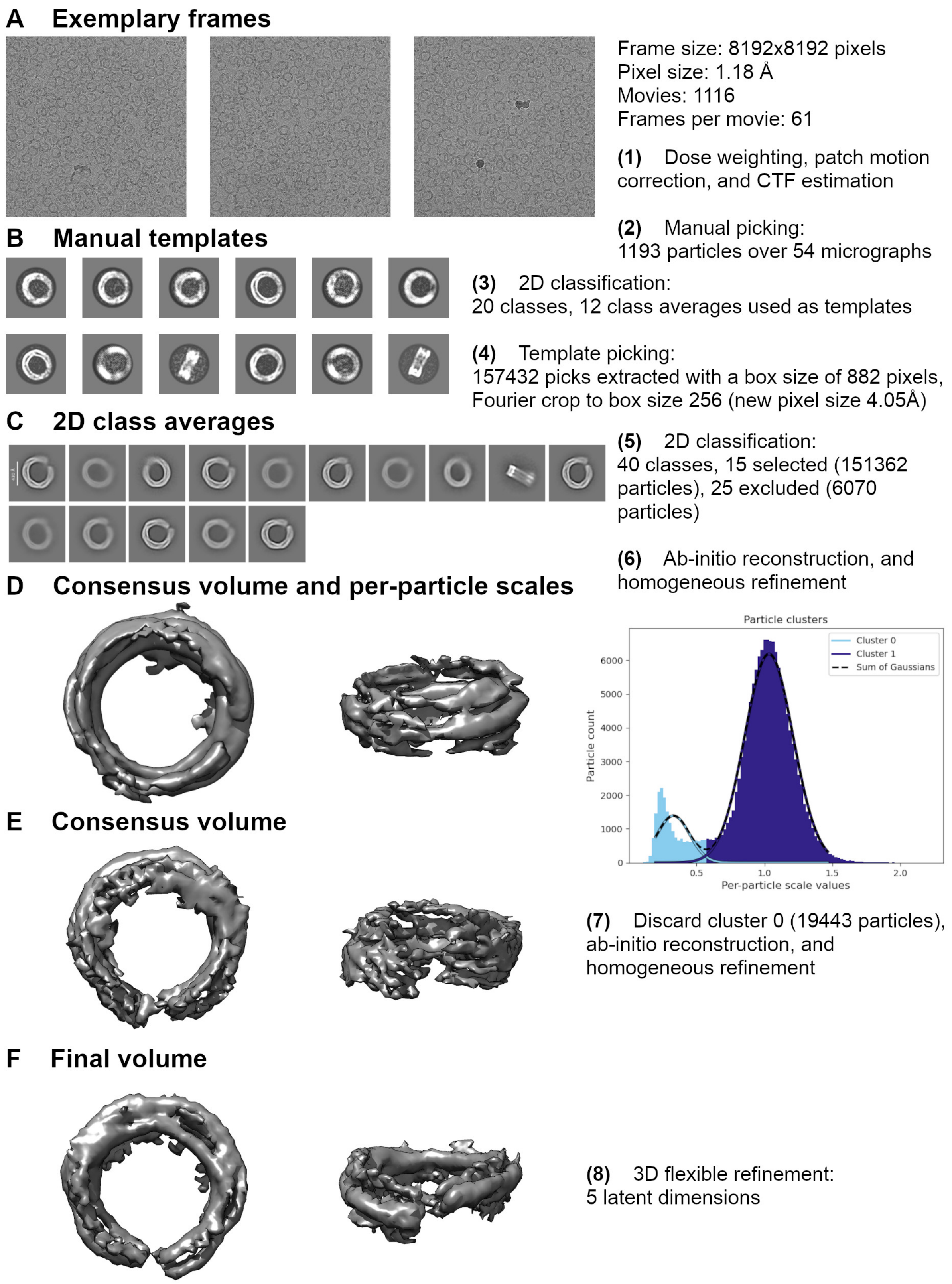


**Supplementary Figure S12:** Detailed single-particle reconstruction process with CryoSPARC v5.0.6. [3] **(A)** Exemplary unprocessed accumulated frames. **(B)** Templates created from 2D class averages after manual particle picking. **(C)** Selected 2D class averages after template picking. **(D)** First consensus volume and per-particle scales, which measure the similarity of observed particle images with the ideal ones given the volume. **(E)** Second consensus volume after discarding particles with low scale values. **(F)** Final volume after 3D flexible refinement.

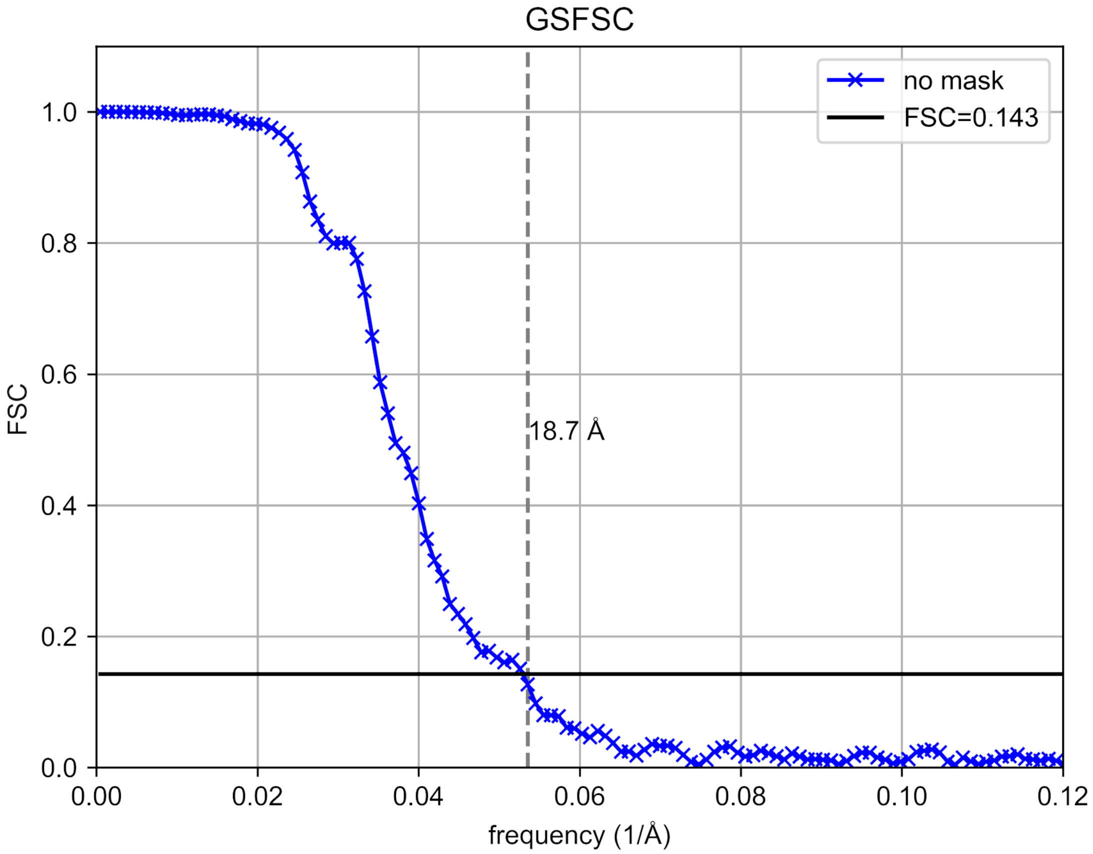


**Supplementary Figure S13:** Gold-standard Fourier shell correlation of the final reconstruction. The blue curve displayes the Fourier Shell Correlation (FSC) values computed between two independent half-volumes. The cutoff value 0.143 is visualized as a horizontal solid black line. The intersection of the FSC curve with the cutoff is emphasized with a vertical dashed grey line. The intersection is located at a spatial frequency that gives the estimated reconstruction resolution as approximately 19 Å.

**Supplementary Equations 1:** Discretization of arc length differences

The method for discretizing arc length differences into base pair counts can be summarized as:

$\forall i \in \{1..N_{\text{bp}}\} : \forall j \in \{1..N_{\text{h}}\}$ :

$$\Delta l_{i,j} = \int_0^{t_i} ||\gamma_i'(t)||\mathrm{d}t + \delta(n_{\text{del},i,j} - n_{\text{ins},i,j}) - \int_0^{t_i} ||\gamma_0'(t)||\mathrm{d}t$$

$\Delta l_{i,j} \geq \delta \implies$ Place an insertion (addition of a base pair) into helix $j$ at index $i$.

$\Delta l_{i,j} \leq -\delta \implies$ Place a deletion (removal of a base pair) into helix $j$ at index $i$.

Here, $N_{\text{bp}} \in \mathbb{N}$ is the total number of base pair positions that will be used while writing the modification pattern into a caDNAno file. $\text{N}_{\text{bp}}$ is solved as the total arc length of the reference curve $\gamma_0$ divided by the value accessible via the "Rise per base pair" field in the settings (default value is 0.34 nm), and rounded to the nearest integer. The position of the reference curve can be specified by the user via the cross section tool included in the graphical user interface. $N_h \in \mathbb{N}$ is the number of helices included in the cross sectional profile as inputted by the user via the cross section tool.

$\Delta l_{i,j}$ is the uncompensated for arc length difference between helix $j$ and the reference curve $\gamma_0$ at index $i$. $\gamma_i$ is the parametric curve ($\gamma_i : [0,1] \mapsto \mathbb{R}^3$) representing the target location for the helical axis of helix $i$. Since $\gamma_i$ is constructed piecewise with natural cubic splines, the integration in the term $\int_0^{t_i} ||\gamma_i'(t)||\mathrm{d}t$ will, in reality, be computed as a sum of integrations over all subintervals contained in $[0, t_i]$. Here, subinterval refers to the closed parameter interval over which the splines used for evaluating the values of $\gamma_i$ remain unchanged. In practice, subintervals will be determined by the placement of target points, specified by the user. The integration in the term $\int_0^{t_i} ||\gamma_0'(t)||\mathrm{d}t$ is performed in a similar fashion.

$\delta$ is the effective length change that follows from the addition or removal of one base pair. This value can be changed via the "Effective rise per modification" field in the settings. $n_{\text{del},i,j}$ and $n_{\text{ins},i,j}$ are the numbers of deletions (base pair removals) and insertions (base pair additions) accumulated in helix $j$ before index $i$. Thus, $\delta(n_{\text{del,j}} - n_{\text{ins,j}})$ acts as a bookkeeping term that ensures that only the previously uncompansated for length differences affect the modification placement at the current index. Primes denote differentiation with respect to $t$, and $|| \cdot ||$ is the $L^2$ (Euclidean) norm.

**Supplementary Equations 2:** Manual modification placement

The number of modifications was determined using the formula [4, 5];

$\Delta l = \theta d$,

where the central angle is $\theta = 2\pi$ for a full revolution, and the interhelical distance is $d = 2.5$ nm.

$\Delta l = 15.707...$ nm $\approx 46$ bp

(Assuming a rise of 0.34 nm along the helical axis per base pair.)

The reference length for the target radius ($R = 20$ nm) is;

$l = \theta R = 125.663...$ nm $\approx 370$ bp

Now, the spacing of modifications can be solved as;

$\frac{\Delta l}{l} \approx \frac{46}{370} \approx \frac{1}{8}$,

since uniform spacing can be applied in the case of constant curvature in one plane.

The cross-sectional profile has three layers of helices perpendicular to the bending direction. Choosing the middle layer as the neutral reference gives the final modification pattern as:

One deletion in each grid cell (8 bp) for helices 1-6.

No modifications in helices 7-12.

One insertion in each grid cell for helices 13-18.

Square lattice twist correction (one deletion in every 63 bp) for all helices.

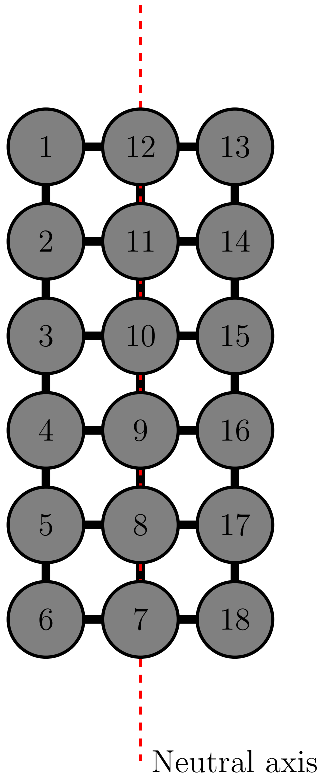

**Supplementary Equations 3:** Parametric equations and sample point sequences for all target curves

$\gamma : [0,1] \mapsto \mathbb{R}^3$

Figure 2 target :

$$\gamma(t) := [R\cos(2\pi t), R\sin(2\pi \text{t}), 0], \qquad R = 20 \text{ nm}$$

$$S := \left\{\gamma\left(\frac{i}{N}\right)\right\}_{i=0}^{N}, \qquad N = 8$$

Figure 3 target :

$$\gamma_1(t) := [R\cos\left(\frac{\pi}{4} + t\frac{7\pi}{4}\right), R\sin\left(\frac{\pi}{4} + t\frac{7\pi}{4}\right), 0], \qquad R = 17 \text{ nm}$$

$$\gamma_2(t) := [2R + R\cos\left(\pi - t\frac{7\pi}{4}\right), R\sin\left(\pi - t\frac{7\pi}{4}\right), 0], \qquad R = 17 \text{ nm}$$

$$S := \left\{\gamma_1\left(\frac{i}{N}\right)\right\}_{i=0}^{N} \cup \left\{\gamma_2\left(\frac{i}{N}\right)\right\}_{i=0}^{N}, \qquad N = 7$$

Figure 4 target :

$$\gamma(t) := [R\cos(4\pi t), R\sin(4\pi t), 2Rt], \qquad R = 21.38 \text{ nm}$$

$$S := \left\{\gamma\left(\frac{i}{N}\right)\right\}_{i=0}^{N}, \qquad N = 21$$

Supplementary Figure S8 target :

$$\gamma(t) := [(R_1 - tR_2)\cos(-4\pi t), (R_1 - tR_2)\sin(-4\pi t), -ht], \quad R_1 = 30 \text{ nm}, R_2 = 20 \text{ nm}, h = 50 \text{ nm}$$

$$S := \left\{\gamma\left(\frac{i}{N}\right)\right\}_{i=0}^{N}, \qquad N = 14$$

Supplementary Figure S9 target :

$$\gamma(t) := [(R_1 + tR_2)\cos(4\pi t), (R_1 + tR_2)\sin(4\pi t), 0], \qquad R_1 = 10 \text{ nm}, R_2 = 40 \text{ nm}$$

$$S := \left\{\gamma\left(\frac{i}{N}\right)\right\}_{i=0}^{N}, \qquad N = 14$$

Figure 5 target :

$$\gamma(t) := [R\cos(2\pi t), R\sin(2\pi t), 0], \qquad R = 21.8 \text{ nm}$$

$$S := \left\{\gamma\left(\frac{i}{N}\right)\right\}_{i=0}^{N}, \qquad N = 8$$

**Supplementary Note 1:** Crossover placement

The crossover placement method implemented by AutoMod can be summarized with simplified pseudocode as:

```
While not all helices within tolerance of target crossover density:
    Find helix i with the smallest crossover density.
    For helices j in the neighbours of helix i, in order of increasing crossover density:
        For index k in shuffled list of all indices:
            If placement conditions:
                Place crossover at index k between helices i and j.
```

In reality, crossovers are categorized into normal crossovers and so-called loops, which are crossovers that cause a helix to have two consecutive crossovers to the same neighbour. The user can select in the settings, whether loops are allowed or not. Even if they are allowed, normal crossovers are preferred by the placement method, since their contribution to overall structural integrity is greater.

The placement conditions contain limitations on how close crossovers can be to the ends of the structure or other crossovers as well as the rotational phases of the participating helices. These conditions can be changed in the user settings via the fields; “No crossovers over last/first”, “Min. crossover separation”, and “Max. phase offset at crossover”. Default values are 6 bp, 6 bp and 3.1 (rad). Even when the maximum phase offset at a crossover is set to a relatively high value, such as the default value, the most well-aligned crossovers are always preferred by the placement method.

Since the above-described method has a minimal amount of conditions restricting the placement of crossovers, several possible crossover patterns exist for most designs. This means that the random shuffling of indices affects the final result. The user can select between different results by changing the “Random seed” field in the settings.

**Supplementary Note 2:** DNAxiS settings

Cross-sectional profile in DNAxiS format [6]:

Helix 1: 324,25,0
Helix 2: 324,22.5,1
Helix 3: 324,20.0,0
Helix 4: 324,17.5,1
Helix 5: 324,15,0
Helix 6: 324,12.5,1
Helix 7: 372,25,1
Helix 8: 372,22.5,0
Helix 9: 372,20.0,1
Helix 10: 372,17.5,0
Helix 11: 372,15.0,1
Helix 12: 372,12.5,0
Helix 13: 420,25,0
Helix 14: 420,22.5,1
Helix 15: 420,20.0,0
Helix 16: 420,17.5,1
Helix 17: 420,15.0,0
Helix 18: 420,12.5,1

Other settings:

$d = 2.5$ nm
crossover factor = 3
alignment threshold = 3
lower bound = 2
same helix crossover spacing = 10
common helix crossover spacing = 7

**Supplementary Note 3:** AutoMod default settings

interhelical gap = 0.5 nm
helix diameter = 2.0 nm
rise per base pair = 0.34 nm
base pairs per turn = 10.5 bp
min. crossover separation = 6 bp
no crossovers over last/first = 6 bp
max. phase offset at crossover = 3.1 (rad)
target crossover density = 0.070 $\frac{1}{\mathrm{bp}}$
margin (over) = 0.005 $\frac{1}{\mathrm{bp}}$
margin (under) = 0.005 $\frac{1}{\mathrm{bp}}$
effective rise per modification = 0.22 nm

**Supplementary Note 4:** Autobreak [2] settings

sequence = M13mp18
readonly = False
rule = xstap.all3
score = sum
func = dG :50
nsol = 20
minlength = 20
maxlength = 60
dontb = 21
verbose = 1
seed = 1
permute = False
writeall = False
csv = True
sort = False

**Supplementary Note 5:** Full oxDNA [7, 8, 9] simulation protocol

Simulation 1: Preliminary relaxation

Simulations for Figures 2 and 5, and Supplementary Figures S1–7 were initiated from neutral conformations where no bending of helices is yet present. Simulations for Figures 3–4, and Supplementary Figures S8–9 were initiated from the target conformation. This was done to avoid the problem of possibly encountering local potential energy minima with the more complex structures.

| | |
|---|---|
| backend = CPU | steps = 2000 |
| sim_type = min | dt = 0.001 |
| verlet_skin = 0.15 | T = 303 K |
| interaction_type = DNA2 | max_backbone_force = 5 |
| salt_concentration = 0.5 | |

Simulation 2: Allowing large deformations to take place

Same as above except:

| | |
|---|---|
| backend = CUDA | sim_type = MD |
| steps = $50 \cdot 10^6$ | (Figures 2 and 5, Supplementary Figures S1–7) |
| steps = $150 \cdot 10^6$ | (Figures 3–4, Supplementary Figures S8–9) |
| thermostat = john | diff_coeff = 2.5 |
| newtonian_steps = 103 | dt = 0.005 |

Simulation 3: Gradually altering backbone potential

Same as above except:

| | |
|---|---|
| steps = $10 \cdot 10^6$ | max_backbone_force = 10 |

Simulation 4: Gradually altering backbone potential

Same as above except:
max_backbone_force = 15

Simulation 5: Gradually altering backbone potential

Same as above except:
max_backbone_force not set

Simulation 6: Trajectory for obtaining the mean conformation

Same as above except:

| | |
|---|---|
| steps = $100 \cdot 10^6$ | print_conf_interval = $1 \cdot 10^6$ |

**Supplementary Data 1:** Staple strand sequences

| Position (caDNAno) | Sequence |
|---|---|
| 14[399] | TTTTTAGCAAGGCCATTTTT |
| 15[77] | TTTCATCAAATCCTAAT |
| 17[292] | TGGTTGCTTACAACATAC |
| 11[243] | GAAAGAGGACTTTCTTAAA |
| 17[114] | GCTGAATATAACTATCATA |
| 13[42] | CAGAGGTGAGAATTATTCAT |
| 17[185] | ATTAAGAGGAAAATGCTTTA |
| 13[250] | GTATTAGACTTCAGATGATG |
| 13[196] | ATTCCCAATTTCATCAGTTG |
| 13[178] | CGGAGATTTGTAGTTAAAGG |
| 11[127] | CAACATGTTTTACGAGGCAT |
| 5[314] | CGCCGCTACATAAAGTGTAA |
| 5[27] | TTTTGGGGTCAGCTGATTGC |
| 7[384] | CCGACCGTGTAATAAGAATA |
| 17[274] | ACGTGGCGAGATGAATCGG |
| 11[190] | GCCGGAACGAGACAACCATC |
| 5[342] | AACGTAACAACTTATGCGATT |
| 17[96] | ACCATCACCCACTGACCTGAA |
| 11[342] | GACAACTCGTCCGTTGTAGCA |
| 13[132] | AAAAGGGCGATCAATCATATGT |
| 15[249] | TGTTCCAGTTTTTATAAATCAA |
| 17[254] | TCATCTTTGACGTCCAATACTG |
| 13[186] | CGTAAAACAAAAGGAATTGAGG |
| 4[210] | CCGGAAACCAGAGGTAAAGATT |
| 17[33] | CCACTATTAAAGTGCGCGAACT |
| 15[282] | ATGCTGTAGCTAGTAAGAGCAA |
| 9[80] | TAATTGCTCCTGGCTTTTGCAA |
| 11[80] | CCACCACACCCGACAATATTAC |
| 17[13] | CGCAACTGTTGGGAAGGGCGAT |
| 17[238] | GCTTTCCTCGTTAGTAATAACATC |
| 16[99] | CGGTCACGCTGCACATTAATTGC |
| 17[316] | GGGCGCGTACTAAGCCGGAAGCA |
| 17[282] | CTGAGAGAGTTGCAGCAAGCGGTC |
| 12[399] | TAGTAAATTGGGCTCAGGCAAGGCA |
| 3[39] | CATCGTAGGAATCATTACCGCGCCGTAG |
| 17[138] | TTGCTCAGAAGTGCCGCATGAGGAAGTTT |
| 0[52] | CGAGGAAACGCAATAATTTCTAAGAACGC |
| 16[153] | AATCGGAACCCTGTTTTTCTTTTCACCAG |
| 10[13] | CGGCTACAGAGGCTTCACCAACCTAAAAC |
| 2[134] | CAGGGTGAAAGGGAGCCCCCGATTTAGAG |
| 3[158] | AGGCGCACAACATTATTACAGGTAGAAAGAT |
| 8[128] | TTGCGGGAGGTTTTGAAAGCAGATAGCCGAA |
| 11[273] | TTTATCAACAATAGATAAGTCCTGAACAAGA |
| 13[118] | TTGGCCTTGAAGTTTGACCATTAGATACATTT |
| 11[16] | GATTAGAGAGTACCTTAAGAAGTTTTGCCAGA |
| 10[283] | TACGTAATGCCACTACGAAGGTGAGGACTAAA |
| 13[344] | GGCATCAATTCTACTACCCTCAGAGCCGCCACC |
| 9[371] | ATAATCAAAATCACCGGAACCAGCCGCCACCCT |
| 3[366] | TAACGTCAAAAATGAAAACACCGGAATCATAATT |
| 16[368] | TCCCCGGGTACCGAGCTTTAAAGGGATTTTAGAC |
| 14[27] | ATTGTAAACGTTAATATTAAATCAGCTCATTTTTT |
| 12[292] | AAAAATCAAAAATCAGGTCTTTTTTATTAAACGGGT |
| 11[12] | AAACGAACTAACGGAATAGGCTGGCTGATCAGACGA |
| 12[117] | GGGGGTAATAGGCGCGAAACAAAGTACAACCGCTTT |
| 17[211] | AGGAGCGGGCGCTAGGGCGCTGGCAAGTGTAGTTG |
| 3[85] | CAGATATAGAAGGCTTATCCGGTAAACGGAATACCCA |
| 6[289] | TATCGGCTGATGCAAATCCACAGTAGGGCTTAATTGAG |
| 10[5] | TACCATTTTTACCCTGACTATTATAGTCAGAAGCAAAGC |

| | |
|---|---|
| 2[98] | AAAGAACTGGCATGATTAAGACTCCTTATTACGCTATCA |
| 2[284] | ACCCACAAAGCGACAGAATCAATCATATATTTTAAATGCA |
| 14[324] | GAAGCCTTTATTTCAATTCGCTATTACGAATTGTTATCCG |
| 12[239] | CCTACATTTTGACGCGAAGAAAGCGAACCTGTCGAATTTA |
| 6[215] | GCGGAACAAAGAAACCACCAGTTTGCCCGAACGTCTGTCCA |
| 4[386] | ACCACATTCAACTAATGCAGATACATAACATAAGGGAACCG |
| 0[113] | GGGATCGTCACCCTCAGCAGCCCGCCACCCTCAGAACCGCCA |
| 13[259] | CTGCGAACGAGTAGATTTTATTCACAAACAAGGTGTACAGACC |
| 10[84] | CTATTAGTTTAGTAGAAAGACAGCATCGGAACGAGGGTAGCAA |
| 15[300] | CGGTGCGGGCCTCGCAAGGATAAAAAGGTCAATAACCTGTTT |
| 14[72] | TCTTTGATTAGAATCAGAGCGCTGTTTCCTGTGTGACCAGCTG |
| 6[336] | TGATAATCAGAAAAGCCCCAAAAAAATTCGCGTCTGGCCAGGCG |
| 1[159] | TTTCAGGGATAGCAAGTAACCGATATATTAATCCGCGACCTGCTC |
| 6[193] | TCTGAAATGGATTATTTATTAAGACGCTGAGAAGAGTCAATAGTG |
| 4[250] | GCACCGTAATCAGTGAATTGAGCGTCTTTCCAGAAATCGCCATATT |
| 13[217] | ATTTAATGGTTTGAAATAAGGTCGACTCTAGAGGATTTTCCCAGTC |
| 9[261] | AAGTACGGTGTCGTCAAATCACCATCCATTAGCAAGGCCGGAAACG |
| 7[366] | GGGTTAGAACCTACCATATCAAAATTATTTGCATCAAAATCATAGG |
| 6[176] | AAGGCCGTAATCATGGTCATGGGAGCTAAACAGGAGGCCGACGAATT |
| 8[62] | AAACCAATCAATAATCGGCTCCATCAAAAATCAGGAAGATTGTATAA |
| 15[344] | ACATTGGCAGATTCACCAGTCACACCCTCAATCAATATCTGGTTTTT |
| 6[118] | AGAGTAAATCGTCGTTTTACATCGGGAGAAACAATAACGGATTCGCC |
| 5[91] | GGGTCAGTGCCTTGAGACCCTCAGAGCCACCACCCTCATCAAAGCGAC |
| 10[147] | CAGCAGCAAATGAAAAATCTAAAGCATCACCTTGCTGAACCT |
| 2[330] | GAAACCATCGATAGCAGCCAACGCTAACGAGTTAAGCCCAATAATAAG |
| 4[399] | CATTATCATTTTTTAAAAGTTTGATTTTTATCAGTGAGGGAAGTGTTTT |
| 16[257] | ATAAATCAATATATGTGAAAAATAATATCCCCATTAAATGTGAGCGAGT |
| 16[178] | AGGATTTAGAAGCAATAACAGCCATAACCAGTATAAAGCCAACGCTCAA |
| 5[301] | AGATCCCAATCCAAATATGCGTTATACAAGCGAAAGGGGGATGTGCTGC |
| 17[369] | GTGAGCTAACTGCGTAACGCCAGCCATTGCATTTAACCTCCGGCTTAGG |
| 7[80] | AGGAACGGTACGCCAGAATCCTGACCACCGAGTAAAAGAT |
| 9[245] | GCGGGGAGAGGCGGCTTGACGGGGAAAGCCGGCGACCAACGCTCTGCCA |
| 8[223] | CCTTCATCAAGAGTAGCCTTTAATTAACGATCTAAAGTGACAGGAGGTTG |
| 11[292] | GTGAATAACCTTGCTTCTTAGAACCCTTAATCAAGTTCCTTCACCGCCTG |
| 1[322] | ATCGCAAGACAAACTCACAATTCCACTGACGAGCACGTATAACGTACTTG |
| 7[298] | GTAATAAGAGAATATAAAATTTTGCACCCAGCTACAAATAGCAATAGCTA |
| 7[98] | TTTCCAGACGTTAGTAAATGAATTTTCTGTATGGGATTTTGCGAATAATA |
| 1[250] | ATTAATTTTCCCTTAGATATACAGTCAAATATCAAACGACCAGTAATAAAA |
| 7[133] | CCTGTAGCATTCCACTTTCGAGGTGAAAGATGAACATAAATCCTCATTAAA |
| 14[236] | AACCAATAAAATCGGCAAAATCCCGGAACAAGAGTGATAGCCCTAAAACAT |
| 16[102] | GCCAACAACCCGTCGGATGAACGGTAATCGTAAAACTAGCATGCATTCAAC |
| 15[30] | AGGCCGGAGACATGGAAGTTTCATTCCATATAACAGTTGAGATTTAGGAAT |
| 3[27] | TGCCAGCTGCATTAAAGGAAGGTCAATCGCAGTTGGAAATAAAGAAATTGA |
| 10[97] | TTTTTATTTTTAGAAAAGTATAGCCCGGAATAGGTGAGTATGTTAGCAAACG |
| 17[80] | GGCGATGGCCCACTACGTGAAGCGTAAGAATACGTGGCGCTGAGACTGATT |
| 5[1] | ACGAACCACCAGCAGCGCCATCATCAAGAAAACAAAATTAATTACTTATCAT |
| 10[340] | CCTATTTCGGAACCAAAGCGAACCAGTGGATAGCCCCCAGCGATTATACCAA |
| 16[204] | AGAACCACCACCATTTTCGGTCATACCTGTAATACTTTTGCGGGATTGGGGC |
| 11[237] | AACTTTCAGCTTGATACCGATAGTTGCGTTTCGTCACCAGTACAAACTACAA |
| 7[311] | GAACGCGAGAAAACTTTTTCAAATATTCCTGATTATTACAAACAATTCATACT |
| 0[23] | TAAAATGTTTAGACACCGGAAGCAAACTCCAATTAATGCCCCCTGCGATTGAG |
| 11[51] | GAAAGAGGCAATGAATCCCCCTCAGCCCGAAAGACTTCAAATATAATATTCA |
| 6[239] | GACTTTTTTCGAGAGGGTTGATTAGCGGGGTTACCACGGAATAAGTTTATTTT |
| 12[108] | ACAGGAAAAACGCTCATGGAAATAAAGGTTATCTAAAATATCTTTAGGAGCAC |
| 3[384] | AATAGCAGCCTTTACTTTTTGCGTTAGATAAATTTTTCAAGCTTGCATGCCTGC |
| 0[318] | ACTAGAAAAAGCCTGTTCATCTTCTGACCTAATCACGCAAATTAAATTAAATCC |
| 8[365] | TCCAAGAACGGAGCAAGCCGATACATAAAGGTGGCAACATATAATTTTTCGGAT |
| 8[27] | GCTCCAAAAGGAATCTTGACAAGAACCGAGTCAGGACGTTAGCTATATTTTCAT |
| 0[117] | TGCCCCTCAGAACCGCCTAACAGTGCCCATTGACGGAAATCTGGAGCAAACAAGA |
| 14[215] | GGAACGGTCTTTCCATTTAACAATTTCATTTGAATTACCTTTTTTAATGGAAACA |
| 16[234] | CGCTCACTGCCCAGCCAGCTTTCCGGCATTTTCGAGCCATCTGAGAGACTACCTT |
| 15[37] | CAATATGATATTCAATCGGCCTCAGGAAGATCGCACTCCGCTTTCCAGTCGGGAA |
| 13[397] | TAAAAATACCTTTTTTAGGGTTGAGTGTAAGAATAGCCCGATTTTTATGAAACAAA |
| 3[280] | AAGTTTTTTCAGAGAGCCACTTTTTAGAATAACATAAAAACAGGGAAGCGCATTAG |
| 12[370] | GCCTAATTTAGATAGTTAGCGTGTATCGGTTTATCAGCTTGCAGACAGCCCTCATA |

| | |
|---|---|
| 14[195] | GATTAAGTTGACCAAAAACATTATGACGCCCCCTTATACTGAACACCCTGAACAA |
| 13[163] | AGCTGATAAATTAGTTTGAGGTAAAGTAATTCTGTCCAGACGACCGTAACCGTGCA |
| 12[330] | CATTATACCGATATTCATTACCCAAATCTTAAGAACTGGCTGCGAGCTGAAAAGGT |
| 11[358] | ACTATTAACATCCAATACTCAGAGCATAAAGCTAAATCGGTTGGGTAACGCCAGGG |
| 10[228] | TGCTTTGAATACCAAGTGCAACAGTGCCACACAGACAATATAAAACCGTCTATCAG |
| 15[348] | TCATTCAGTGAATAAGGCTTGAAAGGAATTGCTAAACAACTTTCAGAACCGCCACC |
| 4[81] | CAGTCTCTGAATTTACCGTTCCAGTAAGCGTCATCATGTACCGTAACA |
| 6[10] | TGGCTTAGAGCTTAATTACCCTCGTTTACCAGACGACGATAAAAACATTGTGTCGA |
| 3[186] | GAGGCGTTTTAGGCAGAACGCGCCTGCGGATTGACCGTATGCCTGAGAGTTATTCC |
| 0[162] | AATCAAGATTAGAGCCCTTTTTAAGAAAAGTAGCCTCGTAGACAGTTGGCAAATCAA |
| 9[38] | ATGCCTGAGTAATGTGTGCAAAGCGCCAGGTGCCTAATGATTGGGTTATATAACAAC |
| 14[272] | ACAATCAATAGAAAATTCAGGCTGAGACTCCTCAAGAGAAGGATTAGTAATAC |
| 2[246] | CGGTCGCTGAGGCTTGCAGGGATCATCGCCTGATAACAAAATAGCGAGATTTGATAA |
| 3[129] | GAGCAAGAAACAATGAATTTTATCCTGAATCTTACCAACATGTAATTTAGGCAGAGG |
| 16[166] | TAACTAATAACAACTAATAGATTAGAGCCGTCAATAGATA |
| 12[367] | AAGATAAAATTCAATTACCTGAGCAAAAGAAGATTTTTCACTCATCGAGAACAGTATT |
| 7[268] | ATGCCTTGCTGGTAATATCCAGACGCGCTTAATGAGCCTGGTTCGCCATTCAGGCTG |
| 5[17] | TTAATCATTGTGAATTACAGCTGCATTTTTTCACGTTGAAAATCTCCAAAAAAAAAGC |
| 7[281] | AAGAATACACACATGAAAGTATTAAGATATGGTTTACCAGCGCCAAAGACACCCCGGT |
| 16[59] | AAAATCCTGTCTCCAACGTCAAAGGGCGATTTTGAATGGCACCGCCTTACAAAATCGC |
| 9[299] | TTACCAGAAGGAAATTAAAGGTGAATTAGGCTATCAGGTCATATGGGATAGGTCACGT |
| 11[62] | CAGGTTTAACGTCAGATGAAATCCTTGAAAAACAACATGTTCAGCTAATCGAACCTCC |
| 2[152] | AGCGTCAGACTGTAGCGATTGAGCGCTAATATCAGAGGCCAGTTACAAAATATCATCA |
| 4[170] | TTTGCGTATTGGGCGCGGTGTAGATGGGCGCATGACAATAACATAGCGATAGCTTAG |
| 14[184] | AAGGAGCGGAATTATCATCATATATTTTAGTTAATTTTAGTATCT |
| 13[152] | GGGAAGAAAAATCTACGTTAATACGCAAATTTTTTAGAACCCGTTTGCCTTTAGGCAGG |
| 12[312] | ACGACGTTGTAAAACGACAATTAAGCAATAAAGCAATCATATGAGATGGTTTAATTTCA |
| 4[362] | TCATTTTTGCGGAGAGATCTACAAATCACCGTCACCGACTTGAGCCTAGCTATTTTTGA |
| 15[379] | ATGCCGGAGAGGGATTTGGGATGATACAGGAGTGTACTGGTAATAAGTTTTAACGGAGG |
| 1[211] | AAACCAAGTACTTTTTCATTAAATTTTTGTTTTGTTAAAATTTTTTTGAAACGCAAAGAC |
| 4[130] | ATTCTTTTATTTTCAGAGGGTACGTTTTCATCGGCAGAGCCGCCGCCAGCATTTTTGTCG |
| 15[237] | GGAGGGAAGGTAAATGTATAAACAGCAGGTCAGGAATCGATTCTCCGTGGGAACAAACGG |
| 3[200] | CCTGAGTAGAAGAACTCAAACATACATTTGATATAATCCTGATTGTTTGGATTTATGTAA |
| 8[11] | TTGCTGTACCGACAAAAGGGGACGACGACAGTACCGTTCTAATCACCAGTAGCACCATTA |
| 8[326] | GTCCAAATATTTAAGGATTGCATCAAAAAGAACAGTTCAGAAAACGAGAATGACCATAA |
| 3[360] | CCGTACTCAGGAGTTCTGAATAAAACACCGGAATCGTCATACGCGTTTTAATTCGAGCTT |
| 9[35] | TTGTTACGGGAGAATTATAGCGTTTGCCATCTTTTCCTCAGAGCCACCAATAGTAGTAGC |
| 6[310] | CACCGCTTCTGGTGCAAAAGGGTGAGAATCACCAATGCCAGAATGGAAAGCGAACTGACC |
| 1[384] | GTACCACGCTGGTTTGCCCCAGCTTCCTGTAGCCAGCTTACGAGCATCAATAGCAAGCAA |
| 7[76] | CAACAGTTAAAGGAACAACTCCCTGACGAGAAACACCAGAATTTTTAGAATAGTCAGCGG |
| 9[106] | CATGTTACTTAGCCCACGCACCCAATAGGAACCACATGGCTTTTGAATTAGAGCCAGCAA |
| 11[179] | ACTGAGCCGACAATGACAGCGCAGACGGTCAATCGCCAAAAGGAATAAATATGCAACTA |
| 1[63] | GCAGAGGCGGCGGTCAGTATTAACTATTAGTCTTTAAAACGTGGATTGATGGTGGTTCCG |
| 4[48] | ACCCTATTATGAGACGGGCAACGAGGTGCCGTAAAGCACTAGGGACATTCTGGCCAACAG |

https://doi.org/10.1063/1.4921957
